# Glutathione supersulphide regulates T-cell receptor signalling

**DOI:** 10.1101/2024.04.30.591985

**Authors:** Yusaku Sasaki, Tadahisa Numakura, Mitsuhiro Yamada, Hisatoshi Sugiura, Tetsuro Matsunaga, Tomoaki Ida, Masanobu Morita, Ayumi Suzuki, Shuichiro Matsumoto, Madoka Kawaguchi, Takeshi Kawabe, Shunichi Tayama, Yuko Okuyama, Tsuyoshi Takata, Kenji Inaba, Satoshi Watanabe, Manami Suzuki, Hirohito Sano, Yorihiko Kyogoku, Rie Tanaka, Ayumi Mitsune, Tomohiro Ichikawa, Naoya Fujino, Tsutomu Tamada, Naoto Ishii, Masakazu Ichinose, Takaaki Akaike, Hozumi Motohashi

## Abstract

Immunometabolism regulates functions and fates of immune cells including T cells. Supersulphides, which are universal metabolites containing catenated sulphur atoms, have various physiological functions based on their unique redox properties. Here we found that activation of T-cell receptor (TCR) signalling was accompanied by supersulphide decrease, which suggests a regulatory contribution of sulphur metabolism to immune function. Consistently, inhibiting supersulphide synthesis facilitated TCR activation and exacerbated allergen-induced type 2 inflammation in mice. Supplementation with glutathione trisulphide (GSSSG), a major endogenous supersulphide, suppressed TCR signalling in naïve CD4^+^ T cells and their differentiation and effectively alleviated the inflammation. Docking simulation revealed interaction of GSSSG with CD3ε chain in the TCR/CD3 complex, which was supported by mass spectrometry detection of persulphidated glutathionylation at a functionally important CXXC motif of CD3ε chain. This study identified a new post-translational modification with supersulfides and demonstrated a critical contribution of sulphur metabolism to TCR signalling regulation.

## Introduction

Recent advances in analytical chemistry have modified our concept of sulphur-containing metabolites^1, 2^. Supersulphides, defined as persulphides and polysulphides containing redox-active sulphur residues with sulphur catenation (S_n_, n > 1)^3^, have been recognized as universal bioactive metabolites that occur in cells and tissues at sub-millimolar levels^1, 2^. Supersulphides exist in cells as low-molecular-weight (LMW) metabolites, such as glutathione hydropersulphide (GSSH), glutathione trisulphide (GSSSG), and cysteine hydropersulphide (CysSSH), and as proteins with persulphidation and polysulphidation at cysteine residue side chains^1,2,3,4,5,6,7,8,9^. Because of an excessive number of sulphur atoms, supersulphides possess a unique chemical property compared with simple thiols: they behave as either nucleophiles or electrophiles depending on the context^1,2,10,11,12,13^. The unique redox property of supersulphides is thought to constitute a molecular basis for their biological roles, such as antioxidant activities^1^, anti-inflammatory functions^8^, protein quality control^10,11^, and mitochondrial energy metabolism^2^.

We previously reported that cysteinyl-tRNA synthetases (CARSs) are one of the major enzymes that synthesize supersulphides in addition to CTH and CBS^2,3^. CARS is a bifunctional enzyme that independently catalyses cysteinyl-tRNA synthesis and CysSSH synthesis. The cytoplasmic isoform CARS1 and the mitochondrial isoform CARS2 are expected to contribute mainly to incorporation of CysSSH into newly synthesized polypeptides and to LMW supersulphide production, respectively. Our clinical study revealed that patients with an asthma-chronic obstructive pulmonary disease (COPD) overlap, which is a refractory type of asthma, showed reduced levels of LMW supersulphides and enhanced inflammatory responses compared with patients with asthma but not COPD, even though both populations of patients were treated with inhaled corticosteroids^14^. Given that asthma is one of the most common allergic diseases in which a pathological CD4^+^ T-cell response is involved, we hypothesized that supersulphide production by CARS2 is required for an adequate CD4^+^ T-cell response.

Activation of naïve CD4^+^ T cells is initiated by antigen stimulation of the TCR/CD3 complex, which activates diverse kinase pathways and transcriptional programs, thereby leading to the production of interleukin (IL)-2 and surface expression of CD25 (IL-2 receptor α)^15,16,17^. Activated T cells undergo rapid clonal expansion and effector T-cell differentiation, in which specific cytokines or combinations of cytokines induce the expression of lineage-specific transcription factors that ultimately guide the development of functionally distinct T helper (Th) cell subsets^15,16,17,18^. Among them, Th type 2 (Th2) cells secrete type 2 cytokines, including IL-4, IL-5, and IL-13, and play fundamental roles in type 2 immune responses, which mediate airway inflammation in asthma^18,19,20^. Type 2 airway inflammation is characterized by elevated IgE levels, eosinophil recruitment, smooth muscle contractility, and mucus hypersecretion^18,19,20^. Activated CD4^+^ T cells undergo broad metabolic reprogramming to initiate clonal expansion and effector differentiation^21,22,23,24,25^. Signalling pathways that control cellular and mitochondrial metabolism, including redox reactions, have crucial roles in T-cell biology^21,22,23,24,25^.

In this study, we first investigated supersulphide levels and *Cars2* expression in naïve CD4^+^ T cells after TCR stimulation. Activation of TCR signalling reduced supersulphide levels and led to reduced *Cars2* expression, which likely reflected altered sulphur metabolism as a part of the metabolic reprogramming during T-cell activation. Consistent with these observations, supersulphide reduction caused by heterozygous deficiency of the *Cars2* gene increased TCR/CD3-mediated early activation of CD4^+^ T cells. As an intriguing result, GSSSG supplementation at a lower dose than its endogenous level limited TCR/CD3-induced activation and differentiation of CD4^+^ T cells. Molecular docking simulation suggested that a unique conformation of GSSSG interacted with the CXXC motif of the CD3ε chain, through which GSSSG likely modulated the assembly and signalling of the TCR/CD3 complex. Indeed, mass spectrometry using Jurkat cell lysates identified persulphidated glutathionylation at the CD3ε CXXC motif only when exogenous GSSSG was added. On the contrary, while the persulphidated glutathionylation was detected on functionally irrelevant cysteine residues with or without exogenous GSSSG. These results have identified persulphidated glutathionylation as a novel protein modification and suggest that the CXXC motif forms a sensitive sensor of supersulphides. The impacts of supersulphides on the regulation of the CD4^+^ T-cell response were verified *in vivo* by using a house dust mite (HDM)-induced type 2 airway inflammation model in *Cars2*^+/–^ mice. This study revealed a new post-translational modification with supersulfides and demonstrated a critical contribution of sulphur metabolism to the regulation of TCR signalling.

## Results

### Supersulphides decrease in TCR signalling

To investigate the contribution of supersulphides to T-cell immunometabolic function, we first studied the effects of TCR signal activation on sulphur metabolism. While cysteine (CysSH) levels were not changed, CysSSH, which is a primary supersulphide synthesized from CysSH, and thiosulphate (HS_2_O_3_^-^), which is an oxidation product of supersulphides, were significantly decreased in naïve CD4^+^ T cells within 3 h after TCR stimulation (Supplementary Fig. 1a). Expression of *Cars2* gene, which encodes one of the major enzymes synthesizing CysSSH from CysSH^2^, was significantly suppressed after 24 h of stimulating naïve CD4^+^ T cells (Supplementary Fig. 1b). These results imply a functional role of supersulphides in regulating TCR signalling in naïve CD4^+^ T cells.

Because supersulphide production by CARS2 is likely to be reduced in activated CD4^+^ T cells, we used *Cars2* mutant mice to evaluate the significance of limited availability of supersulphides for CD4^+^ T-cell activation. Homeostatic T cell development in the thymus and spleen of *Cars2* heterozygous (*Cars2*^+/–^) mice was not different from that of control wild-type (WT) mice (Supplementary Fig. 2a-2f,). We then measured sulphur metabolite levels in naïve CD4^+^ T cells from *Cars2*^+/–^ mice and in WT mice (Supplementary Fig. 2g). We found a significant decrease in supersulphides, including CysSSH, GSSH, and HS O ^-^, in *Cars2*^+/–^ cells compared with WT cells.

### Enhanced TCR signalling by limiting CARS2 function

To clarify the effect of reduced *Cars2* expression on T-cell function, we stimulated naïve CD4^+^ T cells obtained from *Cars2*^+/–^ and WT mice with anti-CD3ε antibody alone or together with anti-CD28 antibody. *Cars2*^+/–^ CD4^+^ T cells expressed relatively high levels of CD25 (Fig. 1a, 1b, and Supplementary Fig. 3a) and CD44 (Supplementary Fig. 3b-3d) after stimulation with anti-CD3ε antibody. This enhanced expression of the activation markers in *Cars2*^+/–^ CD4^+^ T cells was observed after stimulation with 1 μg/ml anti-CD3ε antibody, but not with higher dose (Supplementary Fig. 3e and 3f). CD69 expression was also enhanced in anti-CD3ε antibody-stimulated CD4^+^ T cells from *Cars2*^+/–^ mice (Fig. 1c and 1d). IL-2 secretion was markedly enhanced by heterozygous disruption of *Cars2* (Fig. 1e). These data suggest that CARS2 inhibition significantly augmented TCR/CD3-mediated activation of CD4^+^ T cells.

**Fig. 1:**
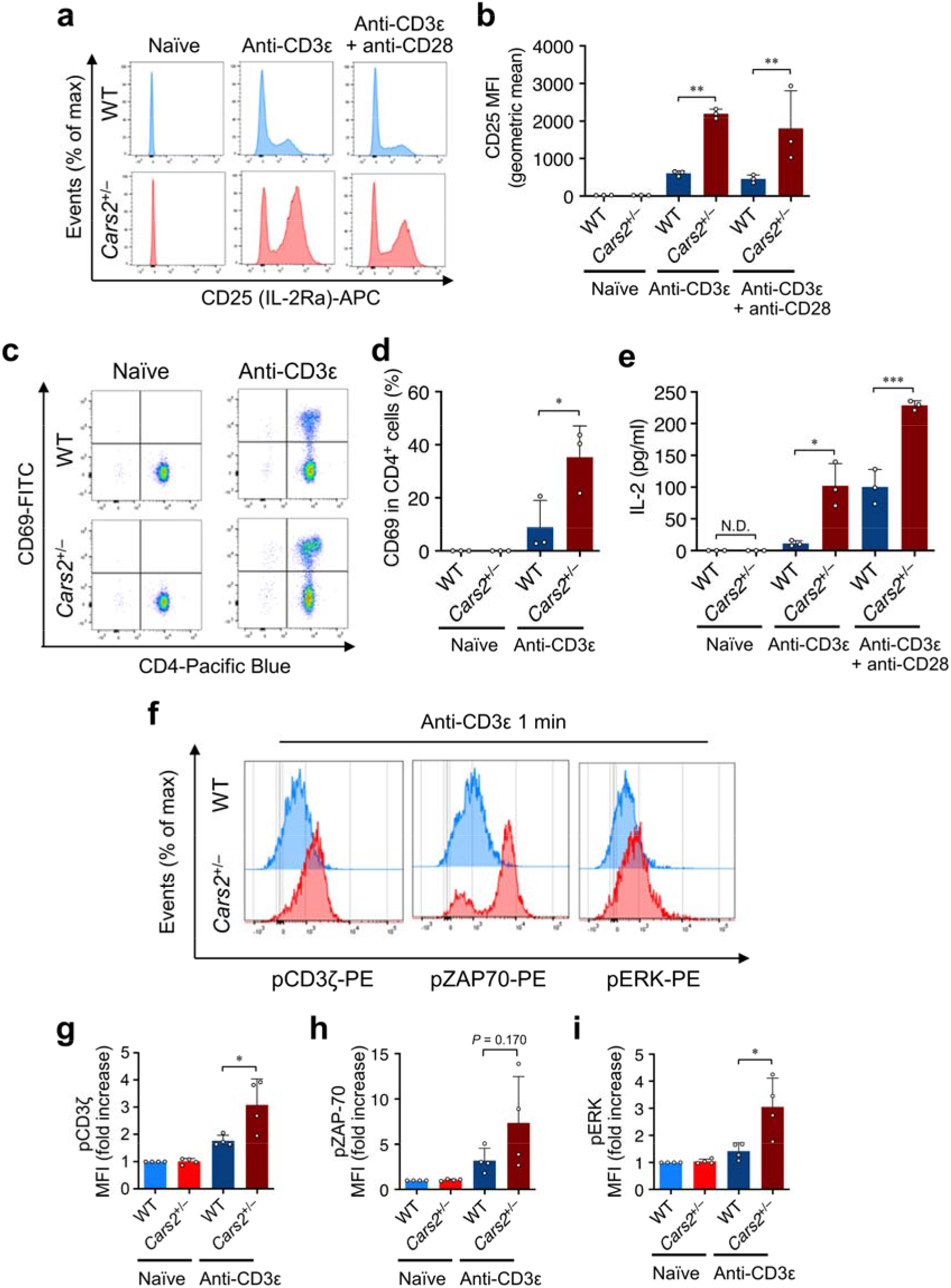
Enhanced CD4^+^ T-cell activation by heterozygous deficiency of *Cars2*. **a-d,** Expression of surface antigens CD25 and CD69 on naïve CD4^+^ T cells obtained from WT and *Cars2*^+/–^ mice as analysed by flow cytometry (FCM) at 24 h after stimulation with anti-CD3ε antibody, or without stimulation, in the presence or absence of anti-CD28 antibody. Representative FCM plots and MFI (mean fluorescence intensity) of surface expression of CD25 on CD4^+^ T cells (**a** and **b**), and representative FCM plots and surface expression of CD69 on CD4^+^ T cells (**c** and **d**). **e,** IL-2 released into the culture medium by activated CD4^+^ T cells obtained from WT and *Cars2*^+/–^ mice analysed via ELISA. **f-i,** FCM analysis of phosphorylation of TCR/CD3 complex components, including CD3ζ, ZAP-70, and ERK, in splenocytes obtained from WT and *Cars2*^+/–^ mice after stimulation with anti-CD3ε antibody. Splenocytes were stimulated with an anti-CD3ε antibody, or were not stimulated, and were then cross-linked with goat anti-Armenian hamster IgG for 1 min. Representative FCM plots of phosphorylation of CD3ζ, ZAP-70, and ERK in CD4^+^ T cells (**f**), and MFIs of pCD3ζ (**g**), pZAP-70 (**h**), and pERK (**i**) relative to that in unstimulated WT cells. Data are mean ± SD. *P* values were analysed with one-way ANOVA with Tukey’s test. \**P* < 0.05; \*\**P* < 0.01, \*\*\**P* < 0.001. Data represent at least two independent experiments.

To investigate the detailed mechanism underlying enhanced activation of *Cars2*^+/–^ naïve CD4^+^ T cells after TCR stimulation, we analysed signalling molecules downstream of the TCR/CD3 complex. Phosphorylation by the Srk family kinase Lck of immunoreceptor tyrosine-based activation motifs (ITAMs) on the cytosolic side of the TCR/CD3 complex occurs early in TCR activation^15^. Lck phosphorylates ITAMs in the TCR/CD3 complex, including the ζ chain^15^. Phosphorylated ITAMs recruit ζ chain-associated protein kinase-70 (ZAP-70). ZAP-70 is then phosphorylated by Lck, which activates ZAP-70 to transduce TCR signalling, thereby inducing T-cell activation^15^. Based on these descriptions, we investigated the phosphorylation of ITAMs and intracellular protein tyrosine kinases in *Cars2*^+/–^ CD4^+^ T cells after TCR stimulation (Fig. 1f-1i, and Supplementary Fig. 4). Compared with WT cells, *Cars2*^+/–^ CD4^+^ T cells showed enhanced phosphorylation of Tyr142 in CD3ζ (Fig. 1f, 1g, Supplementary Fig. 4a and 4d), Tyr319 in ZAP-70 (Fig. 1f, 1h, Supplementary Fig. 4b and 4e), and Thr202 and Tyr204 in extracellular signal-regulated kinase (ERK) (Fig. 1f, 1i, Supplementary Fig. 4c and 4f) after TCR stimulation by anti-CD3ε antibodies. Thus, heterozygous deficiency of the *Cars2* gene enhanced TCR/CD3-mediated phosphorylation of ITAMs, which led to greater phosphorylation of ERK and production of IL-2 in CD4^+^ T cells.

### Inhibition of TCR signalling by GSSSG

Because CARS2 is a mitochondrial bifunctional enzyme possessing two independent activities, supersulphide-synthesizing activity and cysteinyl tRNA-synthesizing activity^2^, CARS2 inhibition should lead to reduced cysteinyl tRNA for mitochondrial translation as well as reduced supersulphide synthesis. To verify that enhanced TCR signalling was attributable to limited supersulphide availability, we supplemented *Cars2*^+/–^ CD4^+^ T cells with one of the major supersulphides *in viv*o, GSSSG, to see whether GSSSG would abolish the aberrantly enhanced TCR signalling with CARS2 inhibition.

GSSSG treatment canceled the enhanced expression of activation markers, including surface antigens CD25 (Fig. 2a and Supplementary Fig. 5a) and CD44 (Fig. 2b and Supplementary Fig. 5b), on anti-CD3ε antibody-treated or anti-CD3ε plus anti-CD28 antibody-treated CD4^+^ T cells from *Cars2*^+/–^ mice. Notably, GSSSG also suppressed the expression of activation markers of CD4^+^ T cells from WT mice. These inhibitory effects on the activation of naïve CD4^+^ T cells were observed under the treatment with 0.1 μM and a higher dose of GSSSG (Supplementary Fig. 5c and 5d). In contrast, glutathione disulphide (GSSG; oxidized glutathione) treatment did not suppress the upregulation of the activation markers induced by CD3 or CD3/28 stimulation on CD4^+^ T cells from both *Cars2*^+/–^ and WT mice (Supplementary Fig. 5e-5h), indicating that the presence of excessive sulphur, or supersulphide, is required for suppressing the TCR signalling. GSSSG treatment significantly reduced anti-CD3ε antibody-induced phosphorylation of CD3ζ (Fig. 2c and 2f), ZAP-70 (Fig. 2d and 2g), and ERK (Fig. 2e and 2h) in CD4^+^ T cells in both *Cars2*^+/–^ and WT mice. These findings collectively indicate that GSSSG reversed the phenotypes of *Cars2*^+/–^ CD4^+^ T cells and suggest that the limited availability of supersulphides is a primary cause of enhanced TCR signalling in CARS2 insufficiency rather than that of cysteinyl tRNA for mitochondrial translation. More importantly, GSSSG also attenuated TCR signalling in WT CD4^+^ T cells, which suggests that supersulphide decrease that occurred in CD4^+^ T cells after TCR/CD3 stimulation is required for full activation of TCR signalling.

**Fig. 2:**
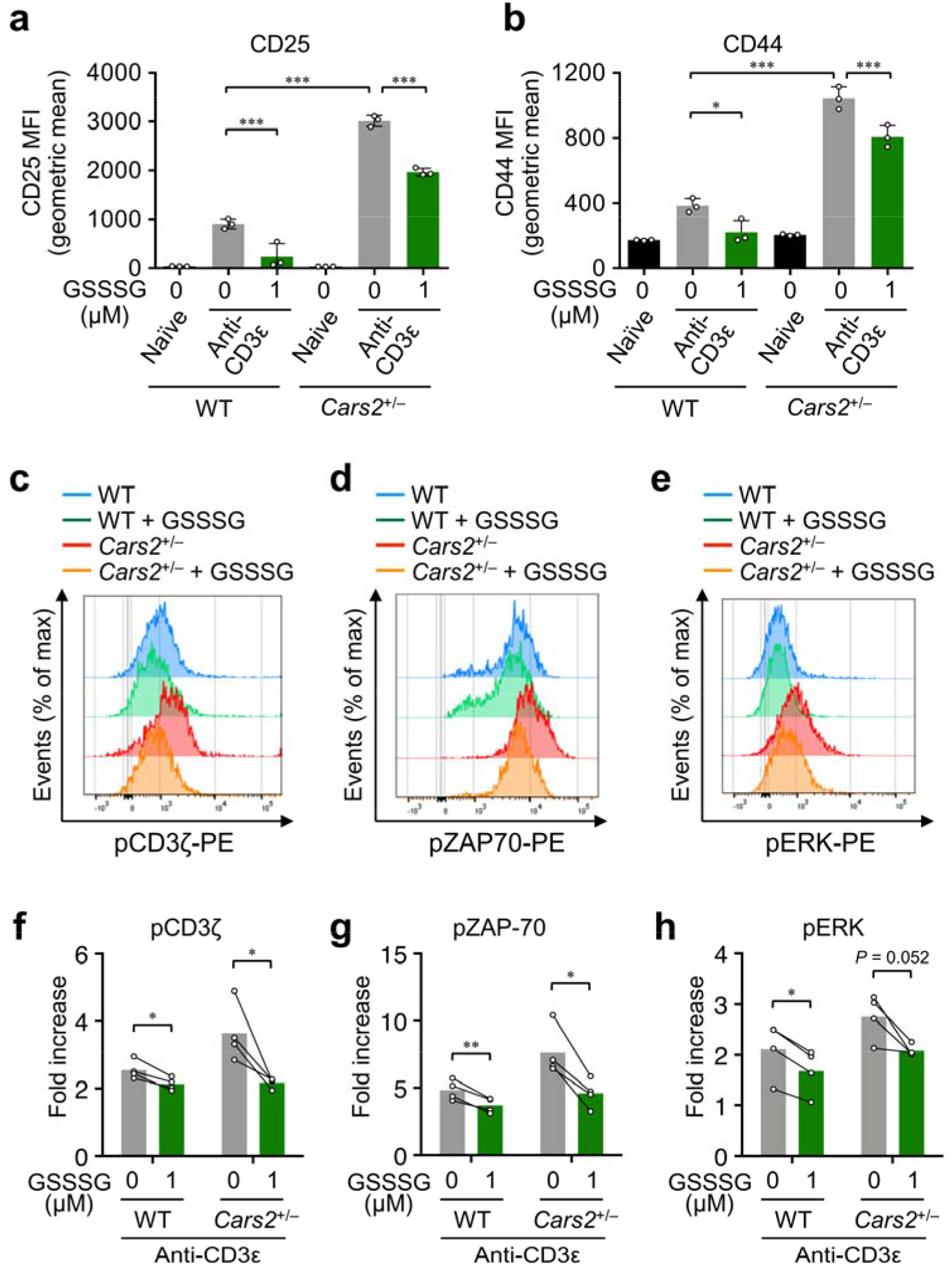
Weakened CD4^+^ T cell activation by GSSSG. **a, b**, Effects of GSSSG on TCR/CD3 signalling-induced expression of surface antigens CD25 (**a**) and CD44 (**b**) as analysed by FCM. Naïve CD4^+^ T cells obtained from WT and *Cars2*^+/–^ mice were stimulated with anti-CD3ε antibody in the presence or absence of GSSSG. Data are mean ± SD. *P* values were analysed with one-way ANOVA with Tukey’s test. \*\*\**P* < 0.001. Data represent at least two independent experiments with consistent results. **c-h,** Effects of GSSSG on TCR/CD3-induced phosphorylation of CD3ζ, ZAP-70, and ERK as analysed by FCM. Splenocytes obtained from WT and *Cars2*^+/–^ mice were treated with GSSSG, or were not treated, for 5 min and were then stimulated with an anti-CD3ε antibody and cross-linked with goat anti-Armenian hamster IgG for 1 min. Representative FCM plots of phosphorylation of CD3ζ (**c**), ZAP-70 (**d**), and ERK (**e**) in CD4^+^ T cells, and MFIs of pCD3ζ (**f**), pZAP-70 (**g**), and pERK (**h**) relative to that in unstimulated WT cells. Data are mean ± SD. *P* values were analysed with a paired *t*-test. \**P* < 0.05. Data were pooled from two independent experiments.

### Inhibition of T cell differentiation by GSSSG

We continued our studies to investigate the impact of heterozygous deletion of *Cars2* on effector CD4^+^ T-cell differentiation as a downstream event of TCR signalling. *Cars2*^+/–^ CD4^+^ T cells cultured under Th1-polarizing conditions had significantly enhanced intracellular expression of T-bet (T-box transcription factor) (Fig. 3a) and type 1 cytokine interferon-γ (IFN-γ) (Fig. 3b), which indicated that CARS2 inhibition enhanced Th1 differentiation. *Cars2*^+/–^ CD4^+^ T cells cultured under Th2-polarizing conditions demonstrated significantly enhanced expression of GATA3, a master Th2 lineage transcription factor (Fig. 3c), and of the type 2 cytokines IL-4, IL-5 and IL-13 (Fig. 3d and Supplementary Fig. 6a-6d). Activated and differentiated T cells from *Cars2*^+/–^ mice proliferated more rapidly than did cells from WT mice (Supplementary Fig. 6e). Expression of the surface activation markers CD25 (Supplementary Fig. 6f-6h) and CD44 (Supplementary Fig. 6i-6k) was also significantly enhanced on activated CD4^+^ T cells from *Cars2*^+/–^ mice compared with cells from WT mice. Dye dilution experiments with CellTrace Violet showed enhanced cell division in activated CD4^+^ T cells from *Cars2*^+/–^ mice compared with cells from WT mice (Supplementary Fig. 6l-6n). These observations—enhanced surface expression of CD25 and CD44 and cell division detected in dye dilution experiments—were also evident in *Cars2*^+/–^ CD4^+^ T cells cultured under Th0 non-polarizing conditions (Supplementary Fig. 6f-6n). As an important result, GSSSG significantly suppressed Th2 differentiation, including expression of GATA3 and IL-13 (Fig. 3e), of CD4^+^ T cells from both *Cars2*^+/–^ and WT mice. Thus, supersulphides suppress TCR/CD3-induced differentiation of CD4^+^ T cells.

**Fig. 3:**
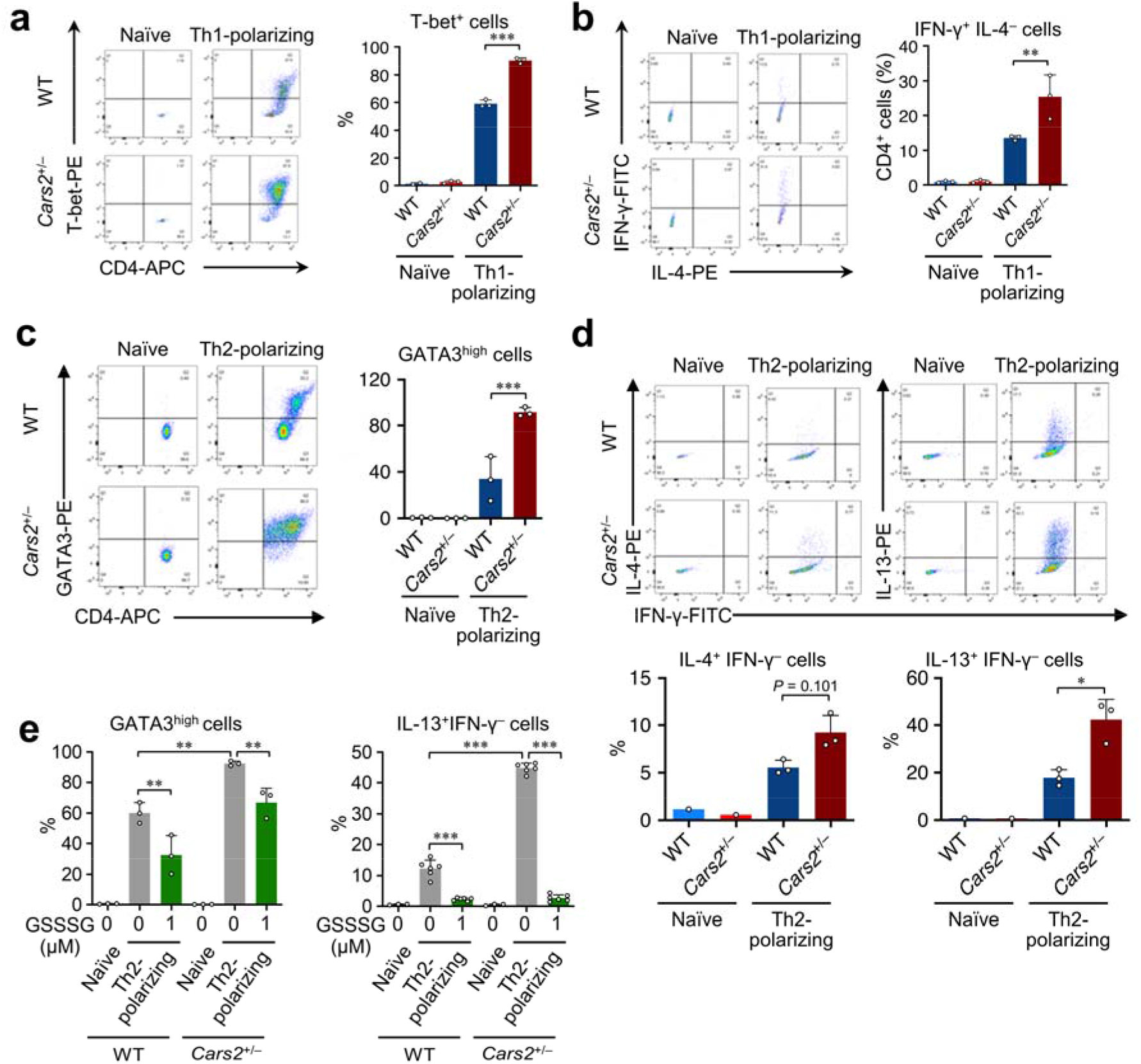
Enhanced T-cell differentiation resulting from heterozygous deficiency of *Cars2 in vitro*. **a, b,** Th1 differentiation of naïve CD4^+^ T cells obtained from WT and *Cars2*^+/–^ mice. Representative FCM plots and intracellular expression of T-bet in CD4^+^ T cells (**a**), and representative FCM plots and intracellular expression of IFN-γ and IL-4 in CD4^+^ T cells (**b**). **c, d,** Th2 differentiation of naïve CD4^+^ T cells obtained from WT and *Cars2*^+/–^ mice. Representative FCM plots and intracellular expression of GATA3 in CD4^+^ T cells (**c**), and representative FCM plots and intracellular expression of IL-4, IL-13, and IFN-γ in CD4^+^ T cells (**d**). **e**, Effects of GSSSG on Th2 differentiation of naïve CD4^+^ T cells obtained from WT and *Cars2*^+/–^ mice. Intracellular expression of GATA3 and IL-13 were analysed by FCM. Data are mean ± SD. *P* values were analysed with one-way ANOVA with Tukey’s test. \**P* < 0.05; \*\**P* < 0.01; \*\*\**P* < 0.001. Data represent at least two independent experiments with consistent results.

### Persulphidated glutathionylation of TCR complex

An important question is how supersulphides control TCR signalling. Because the data above suggested that reduced supersulphide production in naïve CD4^+^ T cells facilitated ITAM phosphorylation, which is an early event in TCR activation, we suspected that the TCR/CD3 complex was a likely target of supersulphides. Moreover, exogenous GSSSG (as low as 0.1 μM) induced maximal inhibitory effects on TCR signalling (Supplementary Fig. 5c and 5d), which was rather surprising in view of the intracellular concentrations of endogenous supersulphides being sub-millimolar^2^. A possible mechanism to explain these data is that exogenously supplied GSSSG serves as a specific ligand for the TCR/CD3 complex.

To investigate the possible interaction between GSSSG and the TCR/CD3 complex, we studied 3D structures of TCRαβ and CD3^26^ and identified a CXXC motif in CD3ε, which is critical for TCR/CD3 complex assembly and signalling^27, 28^, as a candidate acceptor site for GSSSG (Supplementary Fig. 7a). Molecular docking analysis suggested that the sulphur atoms of GSSSG could gain access to the CXXC motif of the CD3ε chain (Supplementary Fig. 7b), by which GSSSG likely modulates the assembly and signalling of the TCR/CD3 complex. We examined whether the CXXC motif in CD3ε is directly modified by GSSSG in the supersulphide proteome analysis for the Jurkat T cell line expressing TCR/CD3 complex (Fig. 4a). A CD3ε peptide containing a CXXC motif (peptide 118-155) was conjugated with glutathione persulphide, forming persulphidated glutathionylation (Supplementary Fig. 7c), only after treatment with GSSSG but not in the basal state (Fig. 4b and 4d). In contrast, persulphidated glutathionylation was detected in a CD3ε peptide without CXXC motifs (peptide 9-64) even in the basal state (Fig. 4c and 4d). These results have identified persulphidated glutathionylation as a novel protein modification and suggest that the CXXC motif is specifically protected from the modification to preserve the responsiveness of TCR signalling. These results also verified that the site-specific increase of persulphidated glutathionylation at the CXXC motif of CD3ε, which is most likely to explain how GSSSG limits the TCR signalling.

**Fig. 4:**
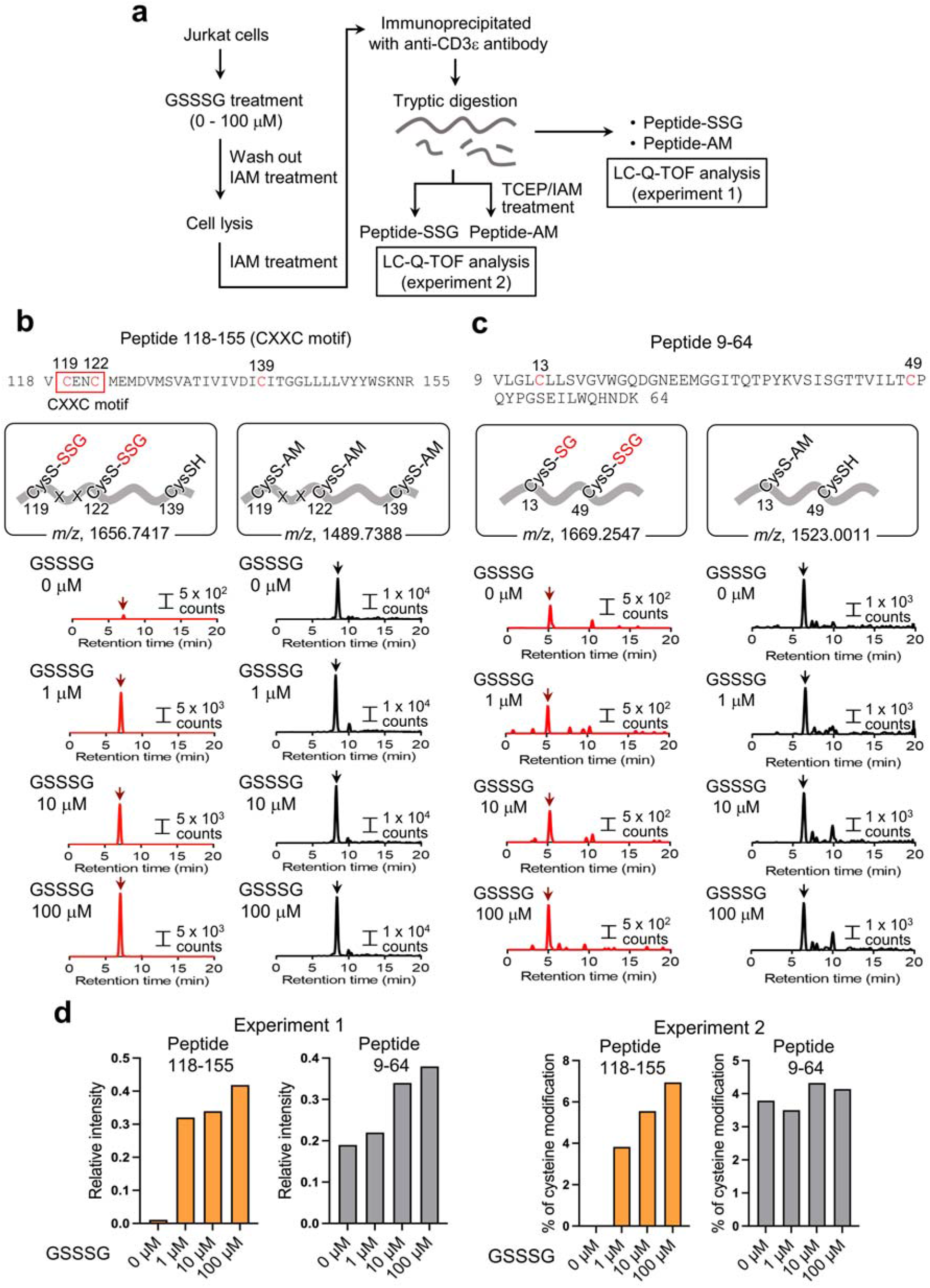
LC-Q-TOF mass chromatograms of a control CD3ε peptide without CXXC motif. **a,** Schematic diagram for identification of modifications in CXXC motif of human CD3ε by the persulphide proteome analysis. **b,** LC-Q-TOF mass chromatograms of a tryptic digestion peptide containing CXXC motif of CD3ε (peptide 118-155) obtained from Jurkat cells treated with or without GSSSG. Left panels shown in red lines are the detection of the CXXC-containing peptide possessing glutathione persulphide adducts (-SSG) at two cystine residues out of three in the peptide, indicating that at least one of the cysteine residues of the CD3ε CXXC motif was involved in the adduct formation. Right panels shown in black lines are the detection of the CXXC-containing peptide not possessing glutathione persulphide adducts but carbamidomethylated cysteine residues (cysteine residues that reacted with IAM; -AM). **c**, LC-Q-TOF mass chromatograms of a tryptic digestion peptide not containing CXXC motif of CD3ε (peptide 9-64) obtained from Jurkat cells treated with or without GSSSG. Left panels shown in red lines are detection of the peptide possessing glutathione and glutathione persulphide adducts at each cysteine residue in the peptide, which are present even in the absence of exogenous GSSSG. Right panels shown in black lines are detection of the peptide not possessing glutathione persulphide adducts but a carbamidomethylated cysteine residue (cysteine residues that reacted with IAM; -AM). **d**, Quantification of peptides with glutathione persulphide adducts. In experiment 1, peptides with glutathione persulphide adducts were normalized to those without. In experiment 2, peptides with glutathione persulphide adducts were normalized to total peptides.

### Severe type 2 airway inflammation asthma in *Cars2*^+/–^ mice

To verify the effect of CARS2 inhibition on TCR signalling in vivo, we subjected *Cars2*^+/–^ mice to intranasal administration of HDM extract, which is an allergen in a murine model of asthma that induces type 2 airway inflammation (Fig. 5a). No significant differences occurred in the number of total cells or cell differentials in bronchoalveolar lavage fluid (BALF) between phosphate-buffered saline (PBS)-treated *Cars2*^+/–^ and WT mice (Fig. 5b). However, *Cars2*^+/–^ mice showed increased eosinophilic inflammation in airways and lungs after HDM treatment compared with WT mice (Fig. 5b-5d). HDM-treated *Cars2*^+/–^ mice had enhanced release of type 2 cytokines, including IL-4, IL-5, and IL-13, in airways compared with HDM-treated WT mice (Fig. 5e). In line with enhanced type 2 inflammatory response in HDM-treated *Cars2*^+/–^ mice, goblet cell hyperplasia in airways of HDM-treated *Cars2*^+/–^ mice increased compared with that in airways of HDM-treated WT mice (Fig. 5f and 5g). Bronchial hyperreactivity (Rrs, resistance of the respiratory system) was also enhanced in HDM-treated *Cars2*^+/–^ mice (Fig. 5h). Moreover, serum IgE levels significantly increased in HDM-treated *Cars2*^+/–^ mice compared with WT mice (Fig. 5i). In lungs of *Cars2*^+/–^ mice treated with HDM, numbers of GATA3^high^ Th2 cells were markedly increased (Fig. 5j and 5k, and Supplementary Fig. 8a-8d), which suggests that CARS2 heterozygous disruption augments allergen-induced type 2 airway inflammation via enhanced activation and Th2 differentiation of naïve CD4^+^ T cells.

**Fig. 5:**
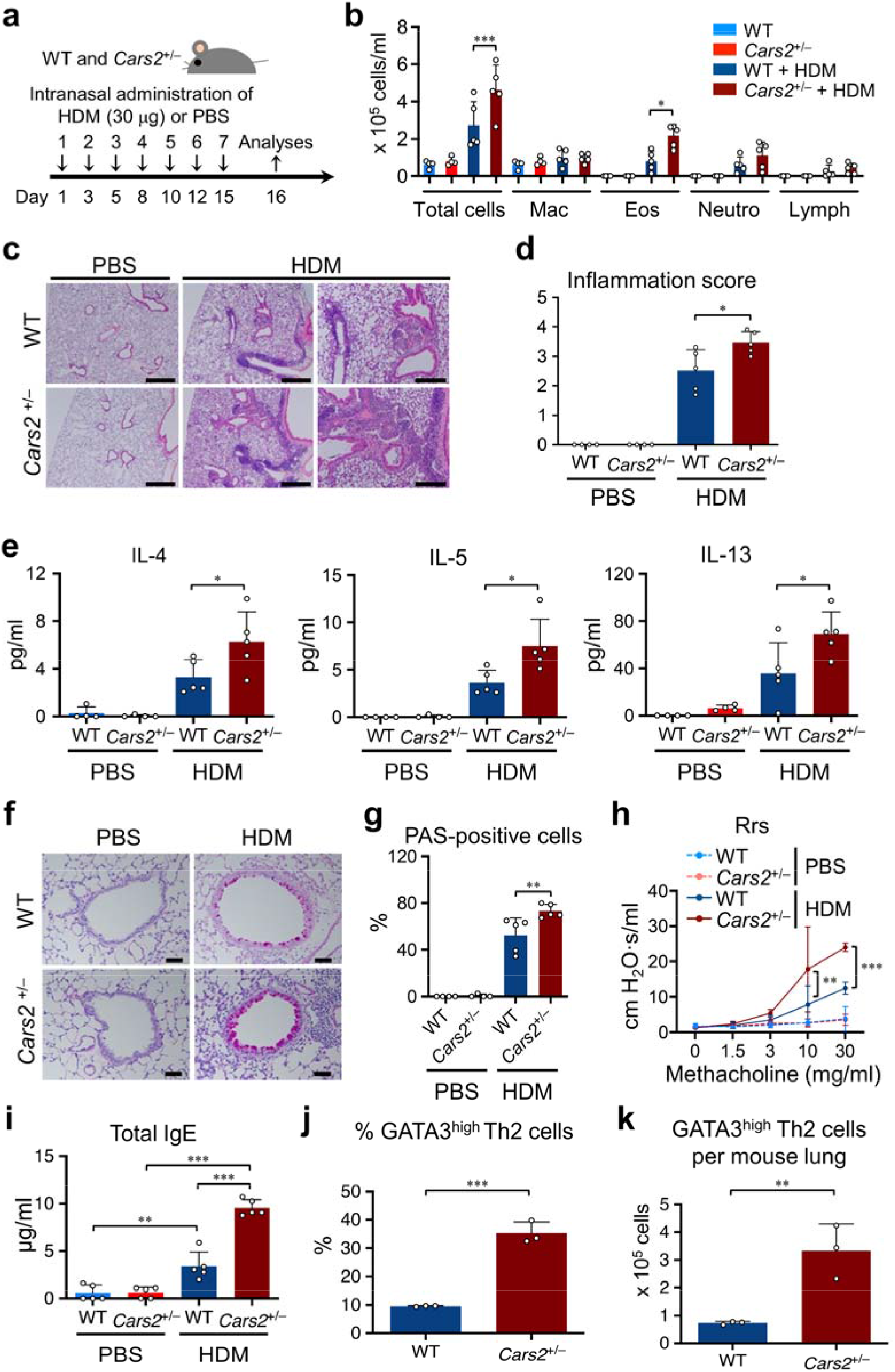
Enhanced allergen-induced asthma in *Cars2*^+/–^ mice. **a,** HDM extract-induced type 2 airway inflammation model. WT and *Cars2*^+/–^ mice were exposed to intranasal HDM or PBS at the indicated times. We obtained and evaluated BALF, blood samples, and lung tissues 24 h after the last HDM exposure. **b,** Numbers of total cells, macrophages (Mac), eosinophils (Eos), neutrophils (Neutro), and lymphocytes (Lymph) in BALF. **c, d**, Representative micrographs of HE-stained lung tissues (**c**) and inflammation score (**d**). Scale bars, 500 µm, left 4 panels, and 200 µm, right 2 panels (**c**). Lung tissue inflammation was semiquantified, and scores were averaged as inflammation scores (**d**). **e,** Levels of type 2 cytokines IL-4, IL-5, and IL-13 in BALF. A cytometric bead array was used for IL-4 and IL-5; ELISA, for IL-13. Each dot represents data from an individual mouse (*n* = 4-5 per group). **f, g**, Representative micrographs of PAS-stained lung tissues (**f**) and PAS-positive cell numbers in airways (**g**). Scale bars, 50 µm (**f**). Each dot represents data from an individual mouse (*n* = 4-5 per group) (**g**). **h**, Respiratory function measured by assessing bronchial hyperreactivity to methacholine. Total respiratory system resistance was measured via a Snapshot-150 perturbation manoeuvre of a flexiVent system. **i**, Total serum IgE measured by ELISA. **j, k,** Percent GATA3^high^ Th2 cells in effector/memory CD3^+^CD4^+^FOXP3^-^CD44^high^ T cells (**j**) and a number of Th2 cells per mouse (CD3^+^CD4^+^FOXP3^-^CD44^high^ GATA3^high^ cells) in lungs in the HDM-induced murine asthma model (**k**). Data are from individual mice (*n* = 4-5 per group, **b, d, e, g**, and **i**; *n* = 3 per group, **h, j** and **k**) and shown as mean ± SD. *P* values were analysed with one-way ANOVA with Tukey’s test (**b, d, e, g,** and **i**), two-way ANOVA with Tukey’s test (**h**), and two-tailed Student’s *t*-test (**j**). \**P* < 0.05; \*\**P* < 0.01; \*\*\**P* < 0.001.

Amounts of IL-33 and thymic stromal lymphopoietin (TSLP), which are epithelium-derived alarmins involved in asthma pathogenesis in that they activate immune cells including type 2 innate lymphoid cells, were not significantly different in HDM-treated *Cars2*^+/–^ and WT mice (Supplementary Fig. 8e and 8f). These data indicate that enhancement of type 2 immune response in HDM-treated *Cars2*^+/–^ mice was independent of the amount of the cytokines, including IL-33 and TSLP, released from injured epithelium in this study.

Both the number of regulatory T cells (Treg) and the percentage of Treg in total lung CD4^+^ T cells in *Cars2*^+/–^ mice at steady state were not significantly different from those in WT mice (Supplementary Fig. 9a-9e). The number of total lung CD4^+^ T cells, Treg as well as conventional FOXP3-negative T cells, in the lungs of *Cars2*^+/-^ mice treated with HDM was higher than that that in WT mice (Supplementary Fig. 9f). However, the percentage of Treg in total lung CD4^+^ cells in *Cars2*^+/–^ mice was not different from that in WT mice (Supplementary Fig. 9g). These data suggest that enhancement of type 2 immune response in HDM-treated *Cars2*^+/–^ mice was not due to the difference in Treg cells between WT and *Cars2*^+/-^ mice.

### Severe type 2 airway inflammation by CARS2 inhibition in T cells

To determine whether CARS2 inhibition in CD4^+^ T cells was responsible for exacerbated type 2 airway inflammation in this model, we next transferred purified CD4^+^ T cells from *Cars2*^+/–^ or WT mice into *Rag2*^-/-^ mice and treated recipient mice with HDM or PBS (Fig. 6a). HDM-treated *Rag2*^-/-^ mice reconstituted with *Cars2*^+/–^ CD4^+^ T cells showed increased accumulation of eosinophils in airways (Fig. 6b) and enhanced release of IL-13, a type 2 cytokine (Fig. 6c). In agreement with these findings, lung inflammation (Fig. 6d) and goblet cell hyperplasia (Fig. 6e) were also enhanced in HDM-treated *Rag2*^-/-^ mice reconstituted with *Cars2*^+/–^ CD4^+^ T cells compared with mice reconstituted with WT CD4^+^ T cells. These data verify that CARS2 inhibition increased the type 2 response through a T-cell-mediated mechanism, which is consistent with our *in vitro* observation of enhanced TCR signalling in *Cars2*^+/–^ CD4^+^ T cells.

**Fig. 6:**
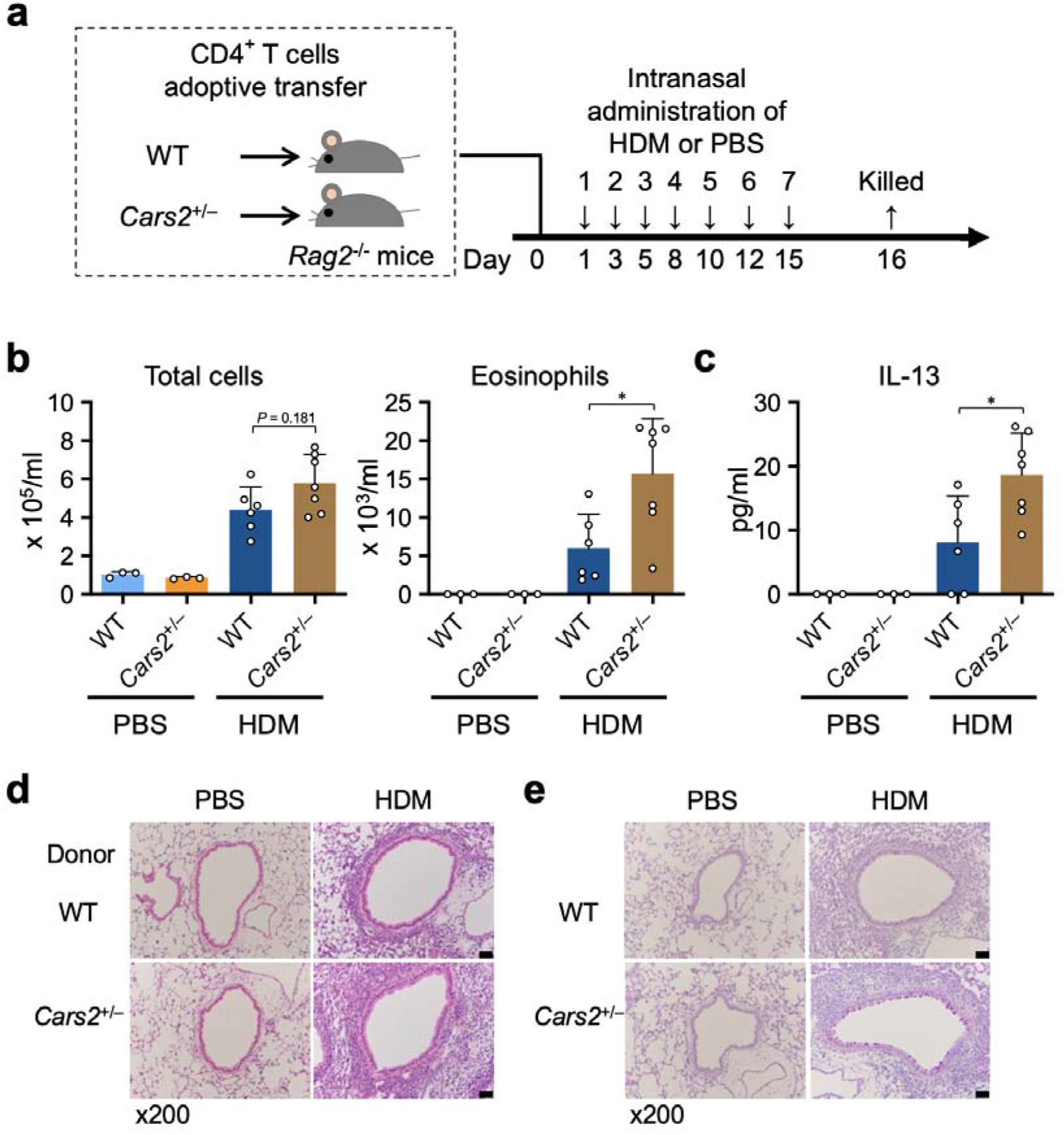
Contribution of CD4^+^ T cells to eosinophilic inflammation in a murine model of asthma. **a,** Schematic illustration of adoptive CD4^+^ T-cell transfer and HDM-induced allergic airway inflammation model. Briefly, splenic CD4^+^ T cells obtained from WT and *Cars2*^+/–^ mice (5 × 10^6^ cells per mouse, *n* = 3-7 per group) were purified by MACS and then intravenously transferred into *Rag2*^-/-^ recipient mice. Recipient mice were then given intranasal HDM or PBS at the indicated times. Twenty-four hours after the last HDM exposure, BALF and lung tissues were obtained for evaluation. **b, c,** Numbers of total cells and eosinophils (**b**) and IL-13 measurements obtained by using ELISA (**c**) in BALF of HDM- or PBS-treated *Rag2*^-/-^ recipient mice after adoptive CD4^+^ T-cell transfer. Each dot represents data from an individual mouse (*n* = 3-7 per group). **d, e**, Representative micrographs of HE-stained (**d**) and PAS-stained (**e**) murine lung tissues. Scale bars, 50 µm. Data are mean ± SD. *P* values were analysed with one-way ANOVA with Tukey’s test. \**P* < 0.05; \*\**P* < 0.01.

### Alleviation of type 2 airway inflammation by GSSSG

We also investigated whether intraperitoneal administration of GSSSG *in vivo* suppressed HDM-induced type 2 airway inflammation (Fig. 7a). Treatment with 1 mM GSSSG significantly suppressed the accumulation of inflammatory cells, especially eosinophils, in the airways of both *Cars2*^+/–^ and WT mice (Fig. 7b). In addition, GSSSG treatment significantly reduced lung inflammation (Fig. 7c) and goblet cell hyperplasia (Fig. 7d) in both HDM-treated *Cars2*^+/–^ and HDM-treated WT mice. Thus, GSSSG has therapeutic potential in CD4^+^ T-cell-mediated type 2 inflammatory disease, including asthma, by shaping TCR signalling (Supplementary Fig. 14).

**Fig. 7:**
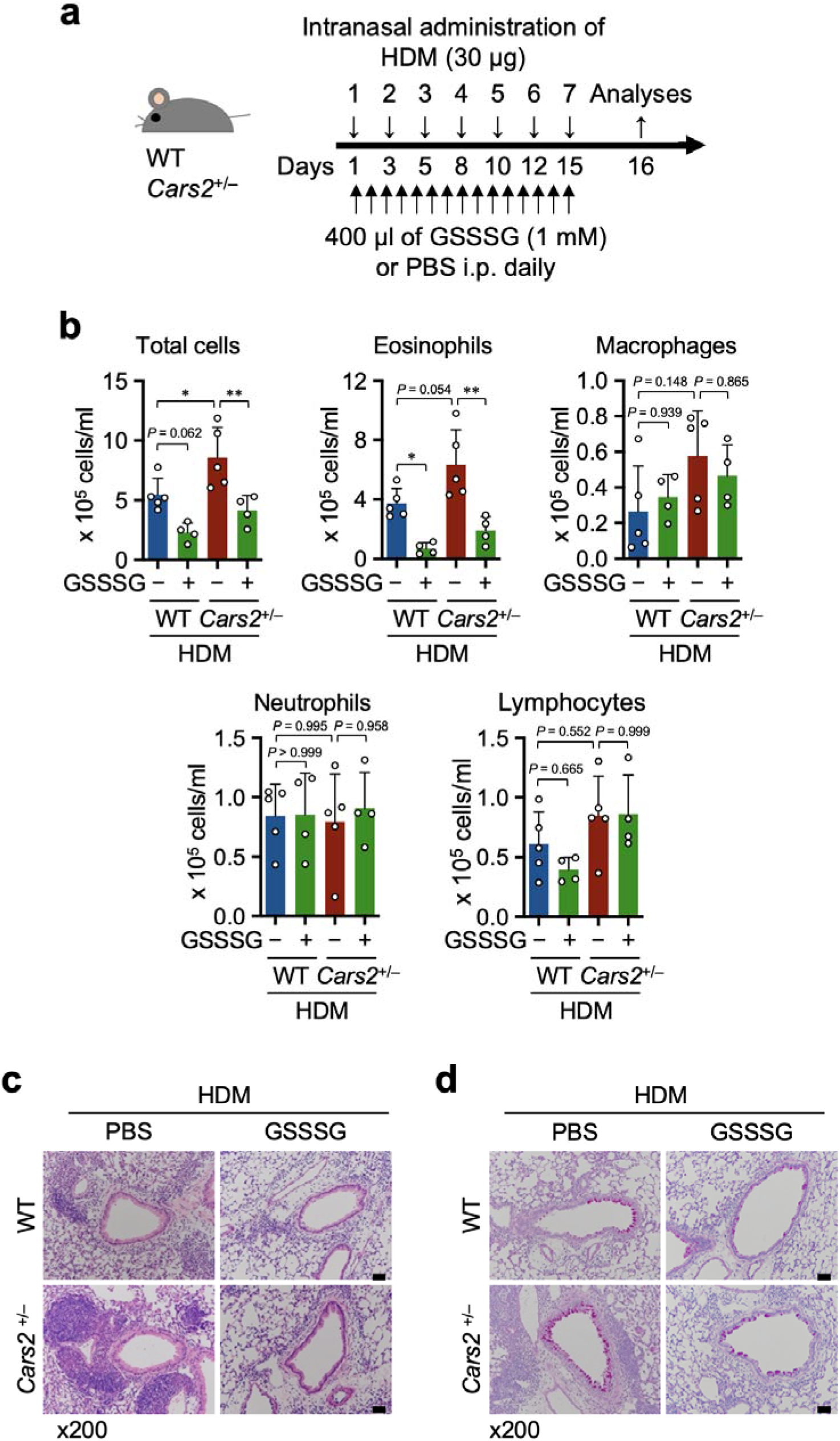
Therapeutic effects of GSSSG on eosinophilic inflammation in a murine model of asthma. **a,** Schematic illustration of the HDM-induced type 2 airway inflammation model treated with GSSSG. Briefly, mice were treated with intranasal HDM with or without daily intraperitoneal (i.p.) GSSSG (1 mM) for 15 days. Twenty-four hours after the last HDM exposure, BALF, lung tissues, and blood were collected for evaluation. **b,** Numbers of total cells, eosinophils, macrophages, neutrophils, and lymphocytes in BALF of HDM-treated mice with or without GSSSG treatment. Each dot represents data from an individual mouse (*n* = 4-5 per group). **c, d,** Representative micrographs of HE-stained (**c**) and PAS-stained (**d**) murine lung tissues. Scale bars, 50 µm. Data are mean ± SD. *P* values were analysed with one-way ANOVA with Tukey’s test. \**P* < 0.05; \*\**P* < 0.01.

## Discussion

TCR-mediated activation and differentiation of T cells are accompanied by reprogramming of cellular metabolism. In this study, we found that TCR signal activation reduced supersulphide levels and led to the suppression of *Cars2* expression in naïve CD4^+^ T cells. Our investigation with *Cars2* mutant mice (*Cars2*^+/–^ mice) revealed that impaired production of supersulphides augmented TCR signalling and worsened allergic inflammation in the HDM-induced murine model of asthma. GSSSG, an abundant supersulphide species *in vivo*, serves as a specific ligand for the TCR/CD3 complex by conjugating the glutathione persulphide residue to the CXXC motif of CD3ε chain, which resulted in attenuation of TCR signalling and consequently alleviation of allergic inflammation *in vivo*. This study elucidated a critical contribution of sulphur metabolism to the modulation of TCR signalling as part of the metabolic reprogramming that regulates T cell activation and revealed that supersulphides function as immunometabolite ligands to generate a newly identified post-translational modification, persulphidated glutathionylation, which mediates suppression of TCR signalling.

An important difference to consider between GSSG (glutathione disulphide) and GSSSG is the electro/nucleophilicity of the sulphur atoms. According to the quantum chemical calculations, both sulphur atoms of disulphide bonds are electrophilic, whereas a central sulphur atom in trisulphide bonds is nucleophilic^3^. Another difference between GSSG and GSSSG is the angle between the two glutathione moieties. Because the three-sulphur-atom chain bends, glutathione moieties get closer to each other and form electrostatic interactions, and the central sulphur atom is positioned at the top front for liganding to the CXXC motif of the CD3ε chain in the TCR/CD3 complex. CXXC motifs, in which two reactive sulphhydryl groups (-SH) are sufficiently close to each other to form disulphides, are important, well-conserved functional elements in proteins^29^. Via reversible oxidation, cysteine residues in the CXXC motif can serve as a regulatory thiol-based switch in proteins and provide an important post-translational control mechanism^30^. Due to the nucleophilicity and structural accessibility of GSSSG, cysteine residues in the CXXC motif that form an intramolecular sulphide bond are likely to recombine to produce intermolecular sulphide bonds with glutathione persulphide residues of GSSSG, resulting in the inhibition of TCR signalling.

Regulation of TCR sensitivity is critical for the survival and homeostatic proliferation of naïve CD4^+^ T cells, which circulate throughout the body and search for antigens. Several factors, such as mTOR (mammalian target of rapamycin) pathway activity, levels of the metabolite adenosine, and signalling via S1P_1_R (sphingosine 1-phosphate 1 receptor), are involved in the maintenance of the size of the naïve CD4^+^ T-cell pool^31^. Unnecessary activation of TCR signalling resulting from dysfunction of these factors ends in an apoptotic loss of naïve CD4^+^ T cells and consequent immune response failure. Alterations in sulphur metabolism causing a substantial decrease in supersulphides in naïve CD4^+^ T cells during TCR activation strongly suggest that supersulphides control and limit TCR sensitivity to prevent abnormal activation of naïve CD4^+^ T cells and thereby preserve the cells.

Although we focused on the direct conjugation of supersulphides to CD3ε chain in this study, alternative targets of supersulphides are also likely involved in the inhibitory effects of supersulphides on T-cell activation and differentiation. Intracellular reactive oxygen species (ROS) constitute one such target. ROS generated by mitochondria or by NADPH oxidase are believed to drive T-cell activation^32^. Supersulphides may suppress TCR responses via quenching ROS that are necessary for full T-cell activation in view of the potent antioxidant capacity. Another possible target of interest is indolamine 2,3-dioxygenase (IDO1). IDO1 is a heme enzyme that oxidizes tryptophan in the first step in the kynurenine pathway, which has an important immunoregulatory function by producing many immunometabolites including kynurenine and serotonin^33^. Because the enzymatic activity of IDO1 is controlled by the redox state of heme iron, i.e., inactive ferric state and active ferrous state, supersulphides contribute to the maintenance of IDO1’s enzymatic activity by reducing ferric heme to ferrous heme. Thus, a reduction in supersulphide levels may leave IDO1 in the inactive state with ferric heme^34^, which would result in a greater T-cell response.

Based on its effective suppression of TCR signalling in naïve CD4^+^ T cells, GSSSG is expected to be a promising therapeutic agent to control immune-mediated diseases, such as asthma and other allergic diseases, which are caused by aberrant activation of CD4^+^ T cells. In this study, we utilized a murine model to assess GSSSG efficacy in immune-mediated diseases. We expect that GSSSG will be effective in controlling type 1 inflammation, in addition to type 2 inflammation, because supersulphide reduction by Cars2 inhibition enhanced Th1 differentiation of naïve CD4^+^ T cells. Therefore, we also believe that developing activators and inhibitors for the supersulphide-synthesizing activity of CARS2, as well as direct administration of GSSSG, will increase the number of possible interventions for various pathological conditions involving T-cell-mediated immunity.

Finally, we must discuss limitations in our study. TCR activation plays a pivotal role in activation of T cells, and our data revealed that sulphur metabolites, which include GSSSG as an immunometabolite ligand for the TCR/CD3 complex, is involved in the regulation of TCR activation. However, it is possible that other relevant receptor pathways including cytokine receptors and their downstream molecules could be regulated by supersulphides because proteins required for T cell subsets differentiation and their functions were shown to possess redox-sensitive cysteine residues that are relevant for their functions^35^. Identification of target proteins of supersulphides would be an interesting issue for future studies to elucidate new regulatory mechanisms of cytokine signalling. Secondly, persulphidated glutathionylation profiles of TCR at different stages of T cell activation and differentiation are still unknown. We speculate that persulphidated glutathionylation of TCR allows fine-tuned regulation of T cell responses to stimuli. Thirdly, although our results revealed that GSSSG covalently binds to a CD3ε CXXC motif, which is critical for TCR-CD3 assembly, structural impacts of the persulphidated glutathionylation on the overall TCR-CD3 complex remain unknown. Finally, how Cars2 expression is downregulated in response to T cell activation remains to be elucidated. Because CARS2-mediated supersulfide synthesis is critical for mitochondrial function^2^, CARS2 inhibition upon T cell activation is likely reasonable considering that activation of TCR signalling switches T cell metabolism towards glycolysis^48^. The same regulatory pathway for the decrease in mitochondrial function may be responsible for the decrease in Cars2 expression.

In summary, this study revealed a pivotal contribution of sulphur metabolism to regulate TCR signalling as a part of the metabolic reprogramming that regulates T-cell activation. The mouse asthma model provided solid *in vivo* evidence for an immunoregulatory function of supersulphides, demonstrating that impaired production of CARS2-mediated supersulphides augments activation and differentiation of T cells and exacerbates type 2 airway inflammation. Our study therefore strongly suggests that sulphur metabolism, particularly supersulphide production, is a new therapeutic target for refractory allergic or immune diseases.

## Methods

### Calculations of supersulphide structures and charge density

Calculations of the optimized structures of all molecules were obtained via the LX 2U Twin 2 server 406 Rh-2 (NEC) with Gaussian 16 (Rev. C.01) at the Research Center for Computational Science. All density functional calculations were investigated at B3LYP/6-311 G(2d,p). We measured charge density by means of natural population analysis with Gaussian 16. We used frequency calculations to confirm no imaginary frequency in optimized structures with the lowest internal energy.

### Materials

Commercially available reagents were obtained as follows: RPMI 1640 medium, fetal calf serum, and penicillin/streptomycin were purchased from Gibco (Invitrogen Life Technologies). 2-β-Mercaptoethanol was purchased from Sigma-Aldrich. HDM extract containing *Dermatophagoides farinae* (10108; 1.08 mg/ml) was purchased from the Institute of Tokyo Environmental Allergy, Inc.. Methacholine chloride was obtained from Tokyo Chemical Industry Co., Ltd..

### GSSSG preparation

We synthesized oxidized GSSSGs as previously described^1, 36^. We diluted GSSSGs in distilled water to prepare a 1 mM stock solution until analysis. Sodium acetate, 30 mM, was added so that the stock solution had a pH of 5.0. For the *in vitro* study, we diluted the stock solution to the proper concentration by using Dulbecco’s Modified Eagle’s Medium (DMEM). For the *in vivo* study, we used the undiluted stock solution.

### Sulphur-omics for murine CD4^+^ T cells

We harvested naïve CD4^+^ T cells from spleens of WT mice and stimulated the cells with anti-CD3ε antibody, or used no stimulation, and we used cells from *Cars2*^+/–^ mice without any stimulation as described above. We homogenized the T cells with a Polytron homogenizer with 70% methanol in 20 mM sodium acetate buffer (pH 6.5) containing 5 mM β-(4-hydroxyphenyl)ethyl iodoacetamide, followed by centrifugation (14,000*g*, 10 min, 4 °C). Supernatants were diluted with 0.1% formic acid containing known amounts of isotope-labelled internal standards, which were then measured with liquid chromatography-electrospray ionization-tandem mass spectrometry (LC-ESI-MS/MS) (LCMS-8060; Shimadzu) coupled to the Nexera UHPLC system (Shimadzu). LC-ESI-MS/MS conditions and synthesis of isotope-labelled internal standards conformed to those given in our previous reports^2^.

We separated supersulphide species by using the Nexera UHPLC and a YMC-Triart C18 column (inner diameter: 50 × 2.0 mm) under elution conditions as follows: mobile phase A (0.1% formic acid) and a linear gradient of 5% to 90% of mobile phase B (0.1% formic acid in methanol) for 15 min with a 0.2 ml/min flow rate at 40 °C. We obtained MS spectra at temperatures of the ESI probe, desolvation line, and heat block of 300 °C, 250 °C, and 400 °C, respectively. We set the nebulizer, heating, and drying nitrogen gas flows to 3, 10, and 10 l/min, respectively. We identified the supersulphide species and quantified them by means of multiple reaction monitoring, as in our previous reports^2^.

### Computational modelling of the 3-D GSSSG-bound TCR/CD3 structure

We employed AutoDock Vina to achieve molecular docking of GSSSG to CD3ε in the TCR/CD3 complex^37^, in which the cryo-EM structure of the TCR/CD3 complex (PDB ID:6JXR) was used. Based on the observation that a CXXC motif is located on the molecular surface only in the CD3ε chain, we set the search area for molecular docking to the vicinity around the CD3ε CXXC motif. We used PyMOL (https://www.pymol.org) to visualize the docking results.

### Detection of persulphidated glutathionylation to CD3**ε**

Immunoprecipitation of human CD3ε from Jurkat cells was performed according to a previous report with slight modifications^38^. Briefly, Jurkat cells (1 x10^7^ cells per tube) were treated with various concentration of GSSSG (0-100 μM) in 2 ml of RPMI 1640 for 30 min at 37°C. The GSSSG-treated cells were centrifuged at 2000 g and washed twice with ice-cold PBS containing 2 mM iodoacetamide (IAM). The cells were lysed in hypotonic buffer (2 mM IAM, 20 mM Tris-HCl pH 7.5) and centrifuged at 15,000 g, and the supernatant was removed. The pellets containing membrane fraction were resuspended and sonicated in 0.5 ml of lysis buffer (1% Triton X-100, 60 mM octyl glucoside, 2 mM IAM, 150 mM NaCl, 20 mM Tris-HCl pH7.5), followed by centrifuged at 15,000 g. The resin pre-reacted with anti-CD3 antibody (Cell Signalling, cat# 85061) and protein G Mag sepharose (Cytiva) was added to supernatants and incubated at RT for 2 hours with rotation. After incubation, immune precipitated samples were washed 5 times with lysis buffer and treated with Trypsin Gold (2 μg/ml; Promega) in 20 mM Tris-HCl (pH7.5) containing 0.04% ProteaseMAX Surfactant (Promega) at 40°C for 2 hrs with shaking. 0.1 ml of 0.1% Formic acid was added to the trypsin-digested samples. After centrifugation (15,000 g, 1 min), the supernatants were subjected to LC-Q-TOF analysis.

The LC-Q-TOF analysis for detecting modifications of cysteine residues of immunoprecipitated CD3ε was performed according to our previous reports with slight modifications^2,3,11,39,40^. Briefly, an AdvanceBio 6545XT LC/Q-TOF (Agilent Technologies) coupled to an Agilent HPLC chip/MS system was used. Tryptic peptides were separated with a microfluidic reversed-phase HPLC chip (ZORBAX 300SB-C18; particle size, 5 μm; inner diameter, 75 μm; and length, 43 mm; Agilent Technologies). Samples were loaded using 0.1% formic acid and 5% acetonitrile in water as a mobile phase at flow rate of 4 μl/min. Peptide separation was carried out using the following elution conditions: 0.1% formic acid in water (mobile phase A) and 0.1% formic acid in acetonitrile (mobile phase B) were used for a linear gradient elution. The flow rate was maintained at 400 nl/min, and the ratio of mobile phase B started at 5% and reached 90% within 15 minutes. The ESI-Q-TOF instrument was set to positive ionization mode as follows; the ionization voltage was 1750 V, and the fragmentor voltage was 175 V at a temperature of 300°C. The mass/charge ratio (*m*/z) values were monitored in the 500- to 1700-Da range with an MS scan rate of 4 s^−1^. Mascot MS/MS ion searches (Matrix Science Web server Mascot version 2.2.) and Agilent MassHunter BioConfirm Software (version10.0) were used to detect and identify the exact *m*/*z* of the peptide containing the carbamidomethylaion (Cys-AM) and other modification [e.g., persulphidated glutathionylation (CysS-SSG), glutathiolation (Cys-SG)]. The modification in VCE NCM EMD VMS VAT IVI VDI CIT GGL LLL VYY WSK NR peptide (peptide 118-155) and VLG LCL LSV GVW GQD GNE EMG GIT QTP YKV SIS GTT VIL TCP QYP GSE ILW QHN DK peptide (peptide 9-64) were detected by monitoring at m/z 1656.7417 and 1669.2547 (for CysS-SSG containing peptide) and m/z 1489.7388 and 1523.0011 (for CysS-AM containing peptide), respectively. In Experiment 1, persulphidated glutathionylation levels were estimated by calculating ratios of CysS-SSG containing peptides against CysS-AM containing peptides. In Experiment 2, to measure total amounts of peptides 118-155 and 9-64, the immunoprecipitated peptides were reduced and re-alkylated in 20 mM Tris-HCl (pH7.5) containing 10 mM TCEP and 50 mM IAM at 40°C for 1 hr. The resultant CysS-AM containing peptides were normalized with ERPPPVPNPDYEPIR peptide (peptide 178-192) (m/z 592.6402) for adjusting digestion efficiency of the protein. The CysS-SSG containing peptides were detected in the same way as in Experiment 1 and similarly normalized with ERPPPVPNPDYEPIR peptide (peptide 178-192) (m/z 592.6402). The persulphidated glutathionylation levels were estimated by calculating ratios of the normalized CysS-SSG containing peptides against the normalized CysS-AM containing peptides.

### Animal studies

We used specific pathogen-free WT and *Cars2*^+/–^ female mice on a C57BL6/J background, 8-12 weeks old^2^. *Rag2*^-/-^ mice were previously described^41^. All mice were kept in specific pathogen-free housing under a constant temperature of 24 °C, the humidity of 40%, and a light cycle of 8:00 A.M. to 8:00 P.M., with food and water available ad libitum.

### HDM-induced murine asthma model

We treated mice under isoflurane anaesthesia with an intranasal application of HDM extracts (indicated dose in PBS) as illustrated in Fig. 5a^42^. Controls were experimental mice treated with PBS. To evaluate the effects of GSSSG in the HDM-induced murine asthma model, we treated mice intraperitoneally with 1 mM GSSSG daily, or no GSSSG, as shown in Fig. 7a.

### BALF cell analysis

Twenty-four hours after the final administration of HDM, we anesthetized the mice and exsanguinated them; lungs were lavaged 3 times with 500 µl of 10 mM PBS. BALF was centrifuged at 800*g* for 5 min, after which supernatants were saved for biochemical analyses, and cell pellets were suspended again in 500 µl of 10 mM PBS. We used a Scepter handheld automated cell counter (Millipore) to count the total cell number. We obtained differential cell counts with cytospin-prepared slides stained with Diff-Quick (Sysmex) or Eosinostain-Torii (Torii Pharmaceutical Co. Ltd.) as described in each figure legend; more than 200 cells were analysed by using conventional morphological criteria.

### Antibodies

For flow cytometry (FCM) analysis, the following antibodies were obtained from BioLegend: anti-CD3ε (145-2C11), anti-CD4 (GK1.5 and RM4-5), anti-CD8a (53-6.7), anti-CD25 (PC61), anti-CD44 (IM7), anti-CD62L (MEL-14), anti-CD69 (H1.2F3), and anti-CD16/32 (93).

Anti-CD4 (RM4-5) and anti-CD69 (H1.2F3) antibodies were purchased from Tonbo Biosciences. For intracellular cytokine and transcription factor staining, fluorochrome-conjugated antibodies including anti-IFN-γ (XMG1.2), anti-IL-4 (11B11), anti-IL-5 (TRFK5), and anti-T-bet (4B10) were obtained from BioLegend. Anti-GATA3 (TWAJ), anti-IL-13 (eBio13A), and anti-Foxp3 (FJK-16s) antibodies were obtained from eBioscience. Phorbol 12-myristate 13-acetate (PMA) and ionomycin were purchased from Merck Millipore. Monensin and brefeldin A were obtained from BD Biosciences. To visualize phosphorylated signalling proteins, surface proteins were stained with anti-CD3ε (eBios500A2) and anti-CD4 (GK1.5) antibodies from eBioscience. Anti-CD247 (CD3ζ) (Tyr142) (3ZBR4S) and anti-pZAP-70/Syk (Tyr319, Tyr342) (n3kobu5) antibodies were purchased from eBioscience. An anti-pERK1/2 (Thr202/Tyr204) antibody (6B8B69) was purchased from BioLegend. Dead cells were stained with the LIVE/DEAD Fixable Near-IR Dead Cell Stain Kit for 633- or 635-nm excitation (Thermo Fisher Scientific), and with 7-Amino-Actinomycin D (7-AAD) Viability Dye (Beckman Coulter). CellTrace Violet was obtained from Thermo Fisher Scientific. For functional studies, anti-CD3ε (145-12C11), anti-CD28 (37.51), and anti-IL-4 (11B11) antibodies were obtained from BioLegend. Anti-IFN-γ antibody used for Th2 differentiation was prepared from R4-6A2 hybridoma (American Type Culture Collection). AffiniPure Goat Anti-Armenian Hamster IgG (H+L) (127-005-160) was obtained from Jackson ImmunoResearch. Recombinant human (rh) IL-2 (200-02), recombinant murine (rm) IL-4 (214-14), and rmIL-12 (210-12) were obtained from PeproTech.

### Isolation of cells from murine lung tissues and FCM

We diced lung tissue samples and digested them with 0.25 mg/ml Liberase (Roche) and 1 mg/ml DNase I (Sigma-Aldrich) at 37 °C for 1 h. Then, cell suspensions and pieces of lungs were mechanically dissociated, passed through a 70-μm cell strainer, and washed with PBS to recover cells. We then purified isolated lung cells by using 2-layer Percoll gradient centrifugation, followed by red blood cell lysis with ammonium-chloride-potassium lysing buffer (Thermo Fisher Scientific)^43, 44^. Single-cell suspensions were blocked with anti-CD16/32 antibodies and stained with fluorochrome-conjugated antibodies against any combination of the following surface antigens: CD3ε, CD4, CD44, CD62L and CD25 (identified above). To study transcription factors, cells were subsequently treated with a Foxp3 Fixation/Permeabilization Kit (eBioscience), according to the manufacturer’s instructions, and were stained with fluorochrome-conjugated antibodies against FOXP3 and GATA3 (identified above) for 30 min on ice.

### Morphometric analysis of lung sections from HDM-induced type 2 inflammation model mice

For histological studies of lung inflammation, we stained lung sections with hematoxylin and eosin (HE). We semiquantified lung inflammation according to the process previously reported with some modifications^45^. Briefly, we chose at least 10 random lung areas, and we graded the severity of perivascular and peribronchial-bronchiolar inflammation as follows: absent, 0; minimal (single scattered leukocytes), 1; mild (aggregates <10 cells thick), 2; moderate (aggregates about 10 cells thick), 3; and severe (many coalescing aggregates thicker than 10 cells), 4. The average calculated points were called average inflammation scores. To quantify goblet cell hyperplasia in the airways, we counted the numbers of periodic acid-Schiff (PAS)-positive cells and total epithelial cells in individual bronchioles. We evaluated at least three medium-sized bronchioles in each slide. We expressed results as the percentage of PAS-positive cells per bronchiole, which we calculated by dividing the number of PAS-positive epithelial cells per bronchiole by the total number of epithelial cells in each bronchiole, as in a previously published report^46^.

### Measurement of respiratory functions in mice

We studied airway resistance at 24 h after the final administration of HDM extract by using the flexiVent system (SCIREQ). Briefly, mice were weighed; deeply anesthetized by means of intraperitoneal injections of medetomidine (ZENOAQ), butorphanol (Meiji Seika Pharma Co., Ltd.), and midazolam (Astellas Pharma Inc.); and tracheostomized. We cannulated the trachea and connected the cannula to the flexiVent system to measure forced oscillations. The mice were ventilated at a tidal volume of 8 ml/kg and respiratory frequency of 150/min with a positive end-expiratory pressure of 2 cm H_2_O, with spontaneous breathing suppressed by an intraperitoneal injection of vecuronium bromide (1 mg/kg of BW) (Fujifilm Wako Pure Chemical Corporation). To assess airway hyperresponsiveness, we administered methacholine chloride (0, 1.5, 3, 10, or 30 mg/ml) (Tokyo Chemical Industry) by means of nebulization. We measured airway resistance by using the Snapshot-150 perturbation manoeuvre method of the flexiVent system. We performed all maneuvers and perturbations until we obtained three correct measurements. For flexiVent perturbations, a 0.95 coefficient of determination was the lower limit for accepting a measurement. For each parameter, we calculated the average of three measurements and show the averages per mouse. The calibration method removed the impedance of the equipment and tracheal tube of this system^42, 47^.

### Measurement of proinflammatory cytokine and chemokine levels

We measured the amounts of cytokines and chemokines, including IL-4 and IL-5, in BALF samples collected from asthma model mice by using a cytometric bead array kit (BD Biosciences) on an LSRFortessa (BD Biosciences) according to the manufacturer’s instructions. We determined the amounts of IL-13, IL-33, and TSLP in the BALF supernatant by means of an enzyme-linked immunosorbent assay (ELISA) and the Quantikine Kit (R&D Systems) according to the manufacturer’s instructions. We determined the amounts of IL-2 in the culture supernatant of activated or naïve CD4^+^ T cells with an ELISA kit (BioLegend). We measured the amounts of IgE in the serum with a commercially available ELISA kit (Sigma-Aldrich).

### Adoptive CD4^+^ T-cell transfer

To reconstitute *Rag2*^-/-^ mice with CD4^+^ T lymphocytes, 5 × 10^6^ CD4^+^ cells were purified from splenocytes obtained from WT and *Cars2*^+/–^ female mice by using murine CD4 (L3T4) Microbeads (130-117-043; Miltenyi Biotec), and cells were transferred into *Rag2*^-/-^ recipients on day 0. Twenty-four hours after CD4^+^ cell transfusion, recipient mice were intranasally treated with HDM as illustrated in Fig. 6a. Twenty-four hours after the last HDM treatment, BALF and lung tissues were collected for analysis.

### Naïve CD4^+^ T-cell isolation and culture

We purified CD4^+^CD25^-^CD44^low^CD62L^high^ naïve T cells obtained from the spleens of WT and *Cars2*^+/–^ littermates with the Naïve CD4^+^ T Cell Isolation Kit II (130-093-227; Miltenyi Biotec) and an autoMACS Pro Cell Separator (Miltenyi Biotech). Cells were cultured in RPMI medium supplemented with penicillin, streptomycin, glutamine, 2-β-mercaptoethanol, and 10% fetal calf serum. For Th2-polarizing conditions, naïve CD4^+^ T cells were plated at a density of 4.0 × 10^5^ cells per ml and stimulated with 1 µg/ml plate-bound anti-CD3ε antibodies (identified above), 1 µg/ml soluble anti-CD28 antibodies (identified above), 5 ng/ml rmIL-4 (identified above), and 1 µg/ml anti-IFN-γ antibodies (identified above). For Th1-polarizing conditions, naïve CD4^+^ T cells were plated at a density of 4.0 × 10^5^ cells per ml and stimulated with 1 µg/ml plate-bound anti-CD3ε antibodies (identified above), 1 µg/ml soluble anti-CD28 antibodies (identified above), 5 ng/ml rmIL-12 (identified above), and 1 µg/ml anti-IL-4 antibodies (identified above). For non-polarizing conditions, naïve CD4^+^ T cells were plated at a density of 4.0 × 10^5^ cells per ml and stimulated with 1 µg/ml plate-bound anti-CD3ε antibodies (identified above), 1 µg/ml soluble anti-CD28 antibodies (identified above), 1 ng/ml rhIL-2 (identified above), and 1 µg/ml anti-IFN-γ antibodies (identified above).

### Analysis of T-cell activation markers

We incubated purified naïve CD4^+^ T lymphocytes (4 × 10^4^ cells) with plate-bound anti-CD3ε antibodies (1 μg/ml) either alone or together with soluble anti-CD28 antibodies (1 μg/ml) for 24 h. For the dose response study, we incubated purified naïve CD4^+^ T lymphocytes (4 × 10^4^ cells) with increasing concentrations of plate-bound anti-CD3ε antibodies (1, 3 or 10 μg/ml) for 24 h. Cells were then washed with PBS containing 1% BSA, blocked with anti-CD16/CD32 antibodies (identified above) for 20 min on ice, washed with PBS containing 1% BSA, and stained to reveal expression of surface markers including CD4, CD25 CD69, and CD44 as indicated. To evaluate the protective effects of GSSSG, naïve CD4^+^ cells purified by using magnetic-activated cell sorting (MACS) were incubated with or without GSSSG (0.1, 1, 10 or 100 µM) together with TCR/CD3 stimulation as described above. Similarly, to evaluate the effects of oxidized glutathione disulphide (GSSG), purified naïve CD4^+^ cells were incubated with or without GSSG (1 µM) together with TCR/CD3 stimulation as described above. Stained cells were identified by using a FACSCanto II flow cytometer (BD Biosciences) and were analysed with FlowJo software (BD Biosciences).

### Analysis of intracellular cytokine production

To stain intracellular cytokines, we cultured MACS-purified naïve CD4^+^ cells under Th2- or Th1-polarizing conditions for 72 h. We then stimulated the cells for 4 h with 100 ng/ml PMA (524400; Merck) and 1 µg/ml ionomycin (407952; Merck) in the presence of GolgiPlug (555029; BD Biosciences) or GolgiStop (552724; BD Biosciences). To assess the effects of GSSSG, we cultured MACS-purified naïve CD4^+^ cells under Th2- or Th1-polarizing conditions with or without 1 µM GSSSG for 72 h. After surface marker staining, cells were fixed and permeabilized with Cytofix/Cytoperm and Perm/Wash Buffer (554714; BD Biosciences) according to the manufacturer’s instructions. Data were acquired by using a FACSCanto II flow cytometer (BD Biosciences) and were analysed with FlowJo software (BD Biosciences).

### Analysis of intracellular transcription factors

To stain intracellular transcription factors, we cultured MACS-purified naïve CD4^+^ cells under Th2- or Th1-polarizing conditions for 72 h. To evaluate the effects of GSSSG, we cultured MACS-purified naïve CD4^+^ cells under Th2- or Th1-polarizing conditions with or without 1 µM GSSSG for 72 h. After surface marker staining, cells were fixed and permeabilized with a Foxp3/Transcription Factor Staining Buffer Set (00-5523-00; eBioscience) according to the manufacturer’s instructions. Data were acquired by using a FACSCanto II flow cytometer (BD Biosciences) and were analysed with FlowJo software (BD Biosciences).

### CellTrace assays

To analyse cell division, we labelled MACS-purified naïve CD4^+^ T cells with 5 μM CellTrace Violet (C34557; Invitrogen Molecular Probes) and then cultured the cells for 72 h in complete RPMI medium (RPMI medium supplemented with 10% FBS, 1% penicillin/streptomycin, and 55 μM 2-β-mercaptoethanol) under appropriate stimulation conditions at 37 °C. Dilution of CellTrace Violet dye was measured by using the FACS Canto II flow cytometer (BD Biosciences), and data were analysed with FlowJo software (BD Biosciences).

### T-cell stimulation for phospho-flow assays

Splenocytes were rested in serum-free medium for 30 min at 37 [, after which they were stimulated with 10 µg/ml anti-CD3ε antibodies (2C11) in the absence or presence of anti-CD28 antibodies (37.51), followed by the addition of 10 ng/ml Goat Anti-Armenian Hamster IgG (127-005-099; Jackson ImmunoResearch Laboratory, Inc.) at 37 [for 1 min. At various times after cross-linking, stimulation was stopped by adding 1 ml of Intracellular (IC) Fixation Buffer (00-8222-49; eBioscience) and incubating the cells at room temperature for 10 min. Cells were stained with the following fluorochrome-conjugated antibodies: allophycocyanin (APC)-conjugated anti-CD4, SuperBright 436-conjugated anti-CD3ε, phycoerythrin (PE)-conjugated anti-phospho CD3ζ, PE-conjugated anti-phospho ZAP-70, and PE-conjugated anti-phospho ERK (identified above). To assess the effects of GSSSG, splenocytes were pre-incubated with or without 1 µM GSSSG before TCR/CD3 stimulation as described above. The stained samples were acquired by using a FACSCanto II flow cytometer (BD Biosciences) and were analysed with FlowJo software (BD Biosciences).

### Quantitative RT-PCR

We used TB Green Premix Ex Taq [(Takara Bio) and a StepOnePlus Real-Time PCR System (Life Technologies) for quantitative RT-PCR. We stimulated naïve CD4^+^ T cells obtained from spleens of WT mice with anti-CD3ε antibody as described above, and total T cell RNA was extracted with a RNeasy Mini Kit (Qiagen). cDNA was then synthesized with the PrimeScript RT Reagent Kit with gDNA Eraser (Takara Bio) according to the manufacturer’s instructions. Each transcript was analysed concurrently on the same plate with analysis of the gene encoding β-actin, and the results are presented relative to β-actin transcript abundance. The primers were as follows: *Cars2* (forward, MA184847-F; reverse, MA184847-R; Takara Bio); *Gapdh* (forward, MA050371-F; reverse, MA050371-R; Takara Bio); and *Actb* (forward, 5′-GGCTGTATTCCCCTCCATCG-3′; reverse, 5′-CCAGTTGGTAACAATGCCATGT-3′).

### Statistics

Data are means[±[SD of at least three independent experiments unless otherwise specified. We assessed comparisons among multiple groups of mice or cells with one-way ANOVA with Tukey’s test, and we used a two-sided Student’s *t*-test for comparisons of two continuous variables unless otherwise specified. *P* values less than 0.05 were taken as significant. We used GraphPad Prism 8 (GraphPad Software, Inc., La Jolla, CA) for comparisons of two continuous variables.

### Study approval

All animal experimental procedures followed the Regulations for Animal Experiments and Related Activities at Tohoku University and the Regulations for Safety Management of Genetic Engineering Experiments in Tohoku University; were reviewed by both the Institutional Laboratory Animal Care and Use Committee of Tohoku University and the Safety Committee for Recombinant DNA Experiments of Tohoku University; and were approved by the President of Tohoku University (#2018MdA-218-02, #2020MdA-139, and #2020MdALMO-115-01).

## Acknowledgements

We thank J.B. Gandy for her editing of the manuscript and evaluating concepts and terminology of the paper with regard to the understanding by non-specialist readers. We thank Mitsu Takahashi at the Department of Respiratory Medicine of Tohoku University Graduate School of Medicine for her excellent technical assistance, and Akihisa Kawajiri at the Department of Microbiology and Immunology of Tohoku University Graduate School of Medicine for his technical advice. Thanks are also due to the Biomedical Research Core of Tohoku University Graduate School of Medicine and the Biomedical Research Unit of Tohoku University Hospital for technical support. This work was supported in part by Grants-in-Aid for Scientific Research [(S), (A), (B), (C), Challenging Exploratory Research, Transformative Research Areas] from the Ministry of Education, Culture, Sports, Science and Technology (MEXT), Japan, to T. Akaike (18H05277, 20K21496 and 21H05263), H. Sugiura (17H04180 and 20H03684), H. Motohashi (20H04832, 21H04799, 21H05258 and 21H05264), M. Yamada (18K08134 and 23K07594), M. Morita (19K07341), T. Ida (20K07306), T. Matsunaga (19K07554), T. Numakura (18K15941 and 21K16106), R. Tanaka (19K17659), Y. Kyogoku (20K17208) and A. Mitsune (21K16131); Japan Science and Technology Agency (JST), CREST Grant Number JPMJCR2024, Japan, to T. Akaike; Japan Agency for Medical Research and Development (AMED) Grant Number JP21zf0127001, Japan, to T. Akaike. This work was also supported by a GSK Japan Research Grant from GlaxoSmithKline K.K. (to R. Tanaka and Y. Kyogoku) and by The Clinical Research Promotion Program for Young Investigators of Tohoku University Hospital (to T. Numakura), and The Basic Research Support Program from Japanese Society of Allergology (to M. Yamada).

## Author contributions

TN, MY and HS conceived the study. YS, TN, MY and HS performed the experiments of T-cell activation, T-cell differentiation, asthma model analysis, and GSSSG treatment in vitro and in vivo experiments. MK performed the experiments of T-cell activation. TK, ST, YO and NI assisted FCM analysis of CD4^+^ T cells. TM, TI, MM and T Takata performed supersulphide metabolome, supersulphide proteome and animal studies. KI and SW analysed the 3D structures of the TCR/CD3 complex bound with GSSSG. AS, SM, MS, HS, YK, RT, AM, NF and TI assisted animal studies and interpretation of in vivo experiments results. T Tamada gave technical advice and interpretation of the results. MI, NI and HS supervised the project. MY, TA and HM designed the study, guided experimental design and analysis. TN, MY, TA and HM wrote the manuscript with input from all authors.

## Additional information

Supplementary Information The online version contains supplementary material. Correspondence and requests for materials should be addressed to Mitsuhiro Yamada, Takaaki Akaike and Hozumi Motohashi. **Reprints and permissions information** is available at www.nature.com/reprints

