## Supplementary information for "Glutathione supersulphide regulates T-cell receptor signalling"

^1^Department of Respiratory Medicine, Tohoku University Graduate School of Medicine, Sendai 980-8574, Japan; ^2^Department of Environmental Medicine and Molecular Toxicology, Tohoku University Graduate School of Medicine, Sendai 980-8575, Japan; ^3^Department of Medical Biochemistry, Tohoku University Graduate School of Medicine, Sendai 980-8575, Japan; ^4^Department of Microbiology and Immunology, Tohoku University Graduate School of Medicine, Sendai 980-8575, Japan; ^5^Institute of Multidisciplinary Research for Advanced Materials, Tohoku University, Sendai 980-8577, Japan; ^6^Medical Institute of Bioregulation , Kyushu University, Fukuoka 812-8582, Japan; ^7^Academic Center of Osaki Citizen Hospital, Osaki 989-6183, Japan; ^8^Department of Gene Expression Regulation, Institute of Development, Aging and Cancer, Tohoku University, Sendai 980-8575.

**^†^**These authors contributed equally to this work.

^‡^These authors should be considered joint senior authors.

***Corresponding authors:**

Mitsuhiro Yamada, Department of Respiratory Medicine, Tohoku University Graduate School of Medicine, Sendai 980-8574, Japan.

Takaaki Akaike, Department of Environmental Medicine and Molecular Toxicology, Tohoku University Graduate School of Medicine, Sendai 980-8575, Japan.

Hozumi Motohashi, Department of Medical Biochemistry, Tohoku University Graduate School of Medicine, Sendai 980-8575, Japan.

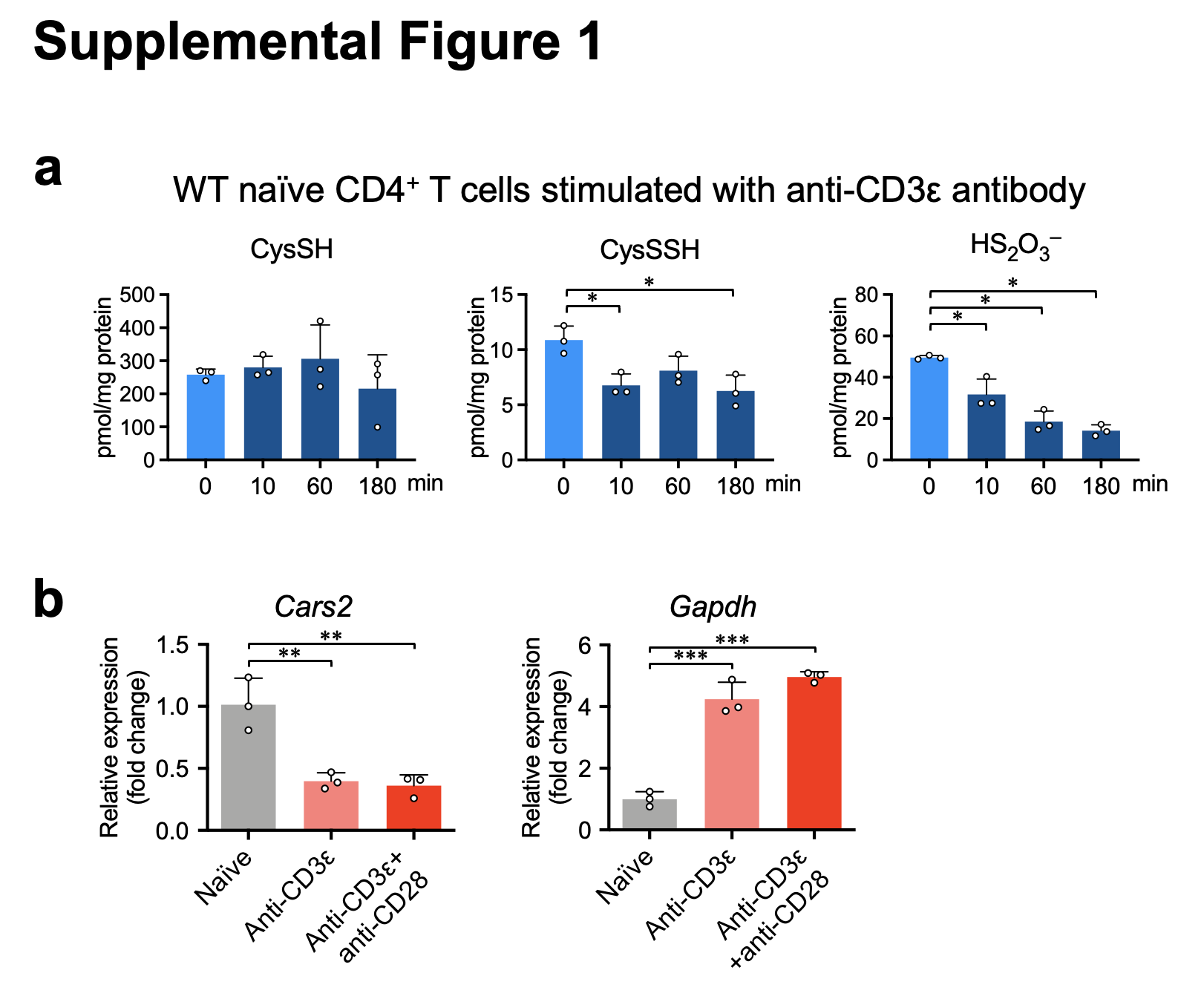

**Supplementary Fig. 1: Sulphur-containing metabolites and *Cars2* expression in CD4^+^ T cells.**

**a,** Sulphur-containing metabolites in naïve CD4^+^ T cells obtained from spleens of WT mice before and after stimulation with anti-CD3ε antibody. Data are mean ± SD (*n* = 3 per group). *P* values were analysed with one-way ANOVA with Tukey’s test. Data represent at least two independent experiments with consistent results. **P* < 0.05.

**b**, Expression of *Cars2* and *Gapdh* in naïve CD4^+^ T cells obtained from WT mice after 24 h of stimulation with anti-CD3ε antibody in the presence or absence of anti-CD28 antibody. Each dot represents data from an individual mouse (*n* = 3 per group). Transcript levels were normalized to those of *Actb* and are presented as relative values to values of unstimulated cells.

Data are mean ± SD. *P* values were analysed with one-way ANOVA with Tukey’s test. Data represent at least two independent experiments with consistent results. ***P* < 0.01; ****P* < 0.001.

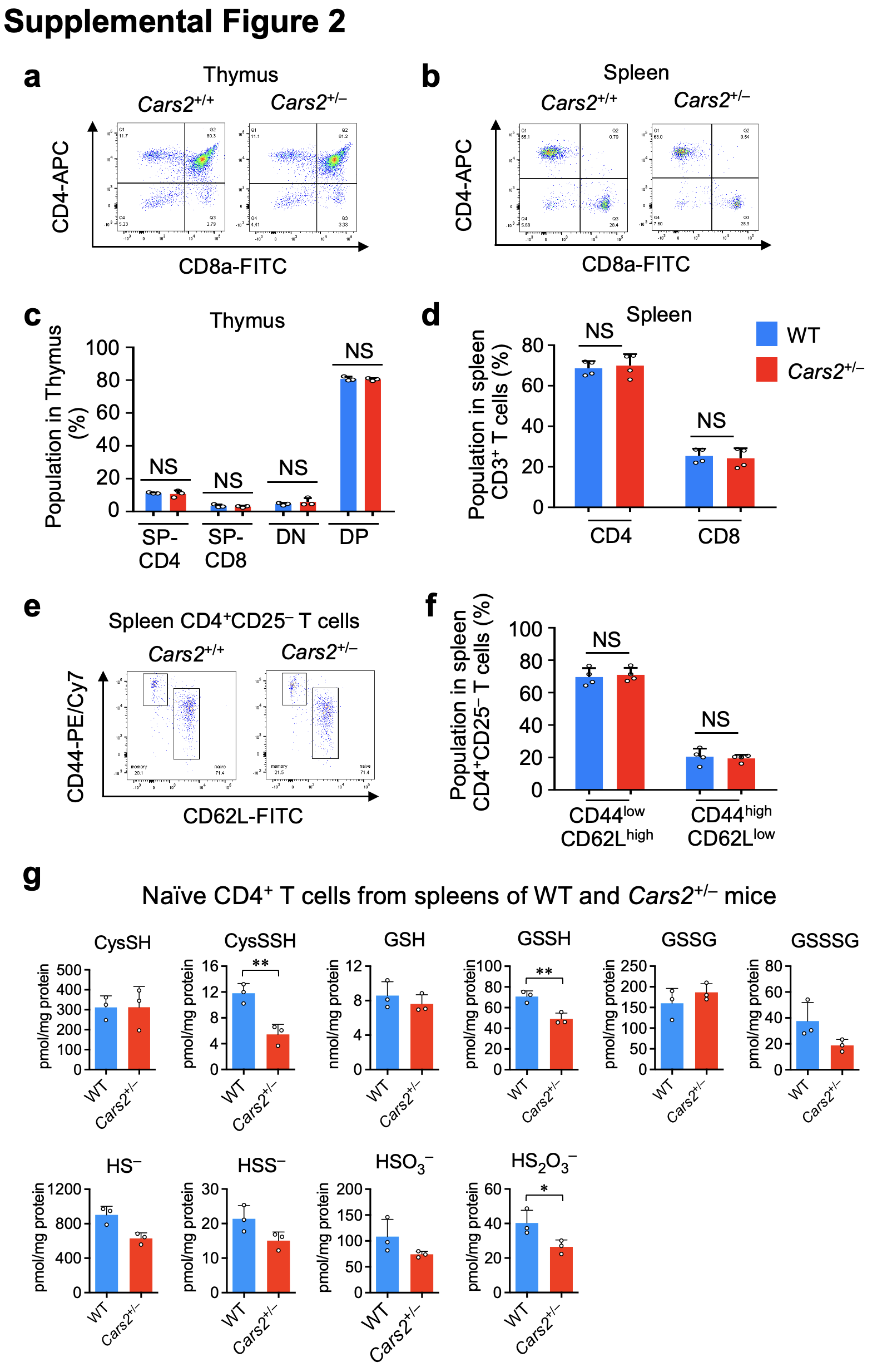

**Supplementary Fig. 2.** **Homeostatic development of T cells is normal in Cars2^+/-^ mice.**

**a, b,** Representative FCM plot depicting the differentiation of thymocytes (**a**) and splenocytes (**b**) from WT and *Cars2*^+/–^ mice into CD4^+^ and CD8^+^ T cells.

**c, d**, Quantification of the differentiated cell types derived from thymocytes (**c**) and splenocytes (**d**) from WT and *Cars2*^+/–^ mice (n = 4 per group). SP-CD4, single positive-CD4; SP-CD8, single positive CD8; DN, double negative; DP, double positive.

**e, f,** Representative FCM plots depicting naïve (CD44^low^CD62L^high^) and effector/memory (CD44^high^CD62L^low^) CD4^+^CD25^-^ T cells in the splenocyte population from WT and *Cars2*^+/–^ mice (**e**) and quantification of each population (**f**). Data from individual mice are shown as the mean ± SD (n = 4 per group). Statistical significance was analysed with two-tailed Student’s t-test. NS; not significant.

**g,** Sulphur-containing metabolites in naïve CD4^+^ T cells obtained from WT and *Cars2*^+/–^ mice.

Data are mean ± SD (*n* = 3 per group). *P* values were analysed with two-tailed Student’s *t*-test. Data represent at least two independent experiments with consistent results. **P* < 0.05; ***P* < 0.01.

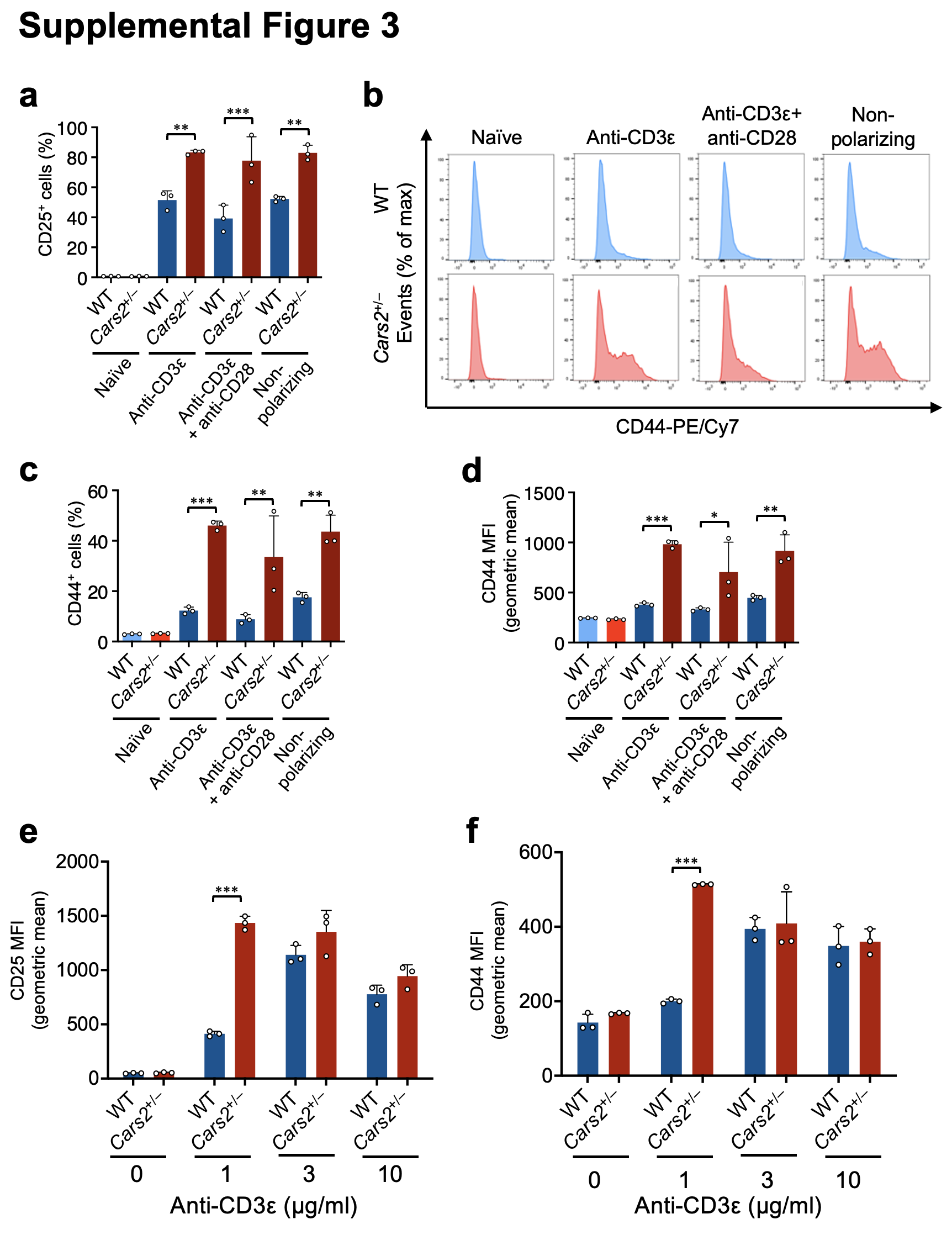

**Supplementary Fig. 3: Enhanced TCR/CD3-mediated activation of naïve CD4^+^ T cells from *Cars2*^+/–^ mice.**

**a-d,** Expression of surface antigens CD44 and CD25 on naïve CD4^+^ T cells obtained from WT and *Cars2*^+/–^ mice analysed by using FCM. Cells were stimulated with 1 μg/ml anti-CD3ε antibody, or were not so stimulated, in the presence or absence of anti-CD28 antibody or anti-CD28 and anti-IFN-γ antibodies and IL-2 (non-polarizing conditions) for 24 h. Representative FCM plots of CD44 (**b**) and surface expression of CD25 (**a**) and CD44 (**c** and **d**) on CD4^+^ T cells. PE/Cy7, phycoerythrin-cyanine 7 (a surface-labelling fluorochrome).

**e, f,** Expression of surface antigens CD25 (**e**) and CD44 (**f**) on naïve CD4^+^ T cells after stimulation with different indicated doses of anti-CD3ε antibodies.

Data are mean ± SD. *P* values were analysed with one-way ANOVA with Tukey’s test (**a** and **c**-**f**). Data represent at least three independent experiments with consistent results. **P* < 0.05; ***P* < 0.01; ****P* < 0.001.

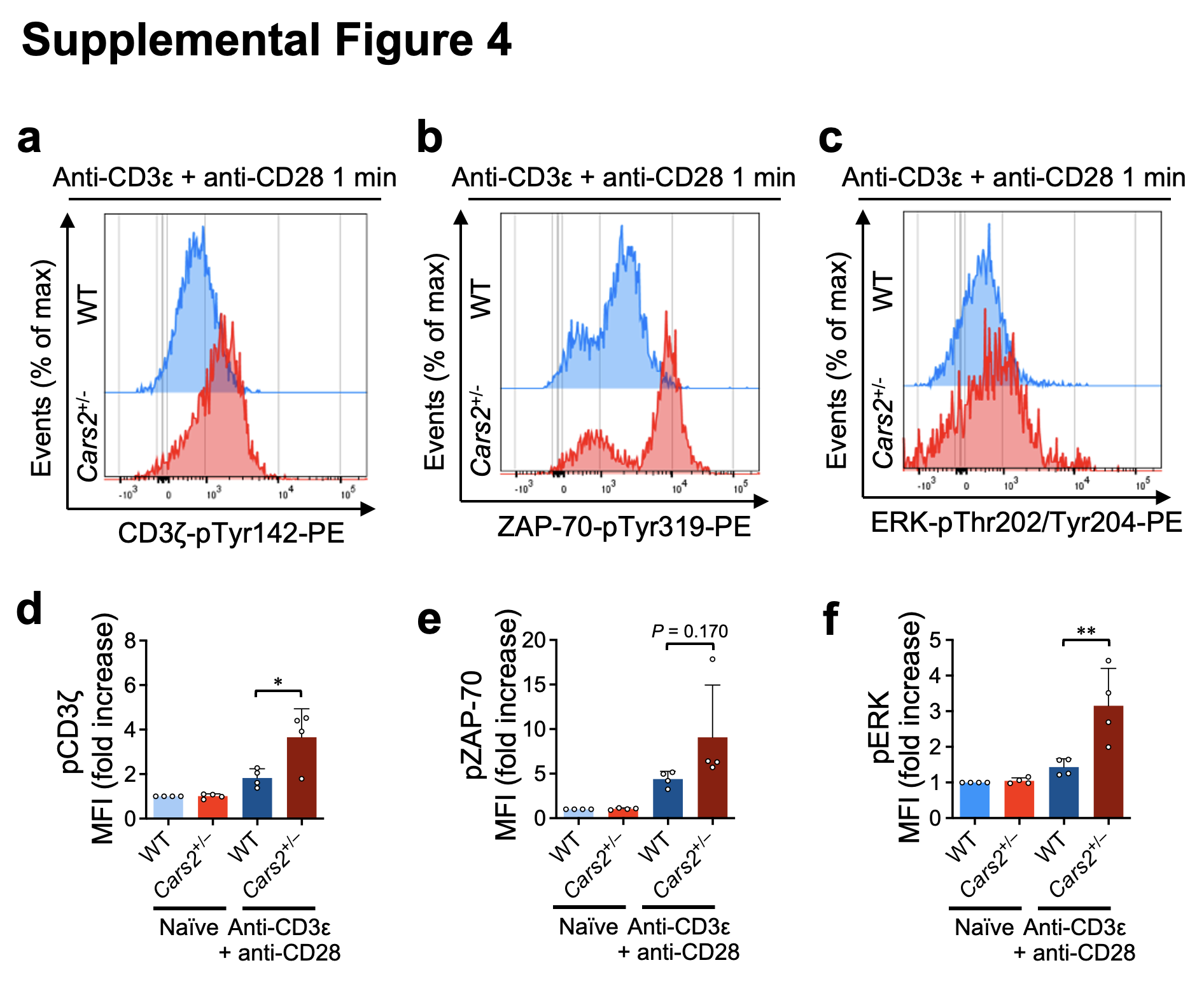

**Supplementary Fig. 4: FCM analysis of phosphorylation of CD3ζ, ZAP-70, and ERK after TCR/CD3 stimulation with CD28-mediated co-stimulation.**

**a-f,** Splenocytes obtained from WT and *Cars2*^+/–^ mice were incubated with anti-CD3ε and anti-CD28 antibodies, or were not so incubated, and were then cross-linked with goat anti-Armenian hamster IgG for 1 min. Representative FCM plots of phosphorylated CD3ζ (**a**), ZAP-70 (**b**), and ERK (**c**) in single/CD3^+^/CD4^+^ gates. Mean fluorescence intensity (MFI) of pCD3ζ (**d**), pZAP-70 (**e**), and pERK (**f**) relative to MFIs of the same proteins in unstimulated WT cells. Data are mean ± SD. *P* values were analysed with one-way ANOVA with Tukey’s test (**d-f**).

Data represent at least three independent experiments with consistent results. **P* < 0.05; ***P* < 0.01.

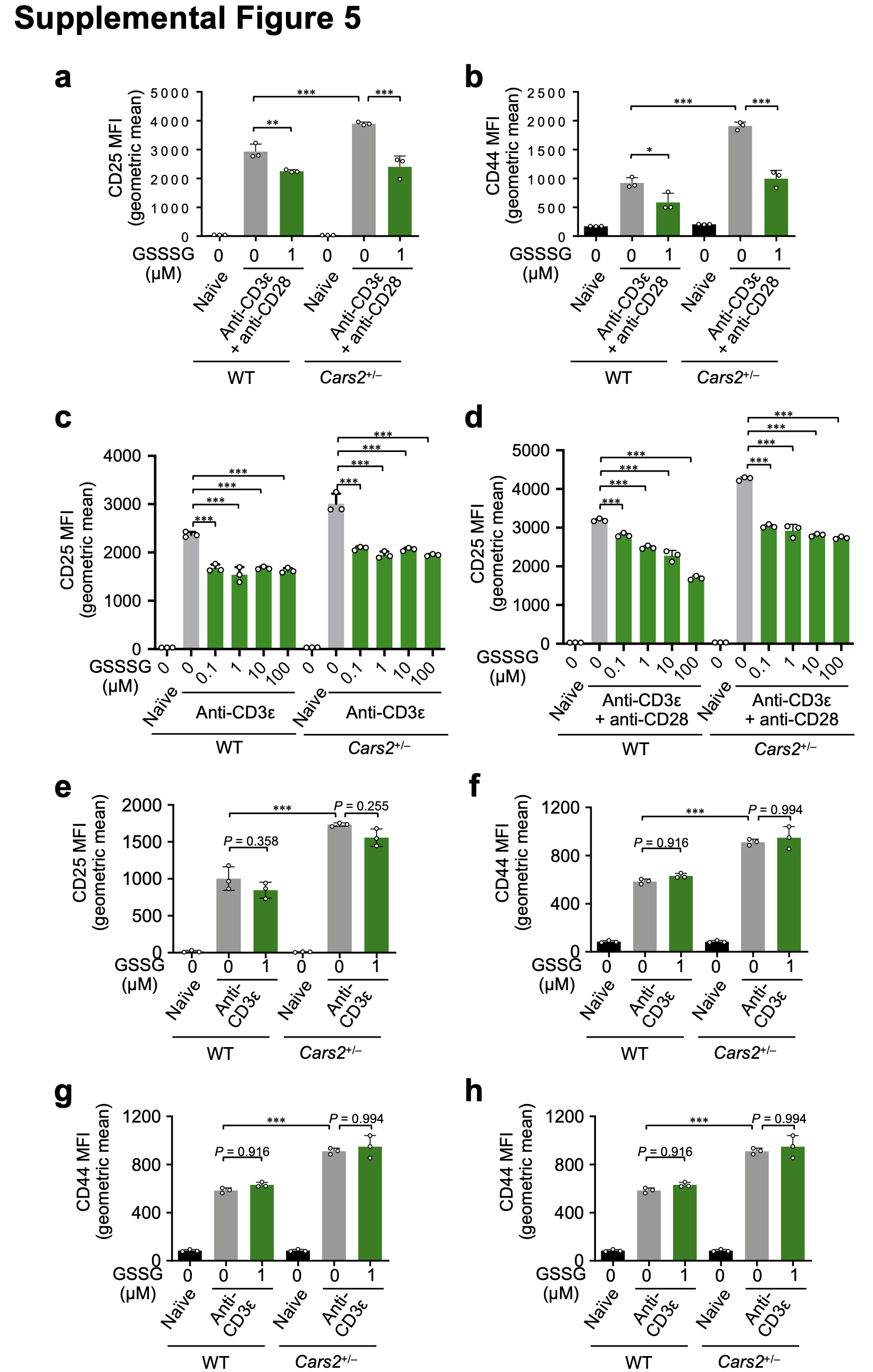

**Supplementary Fig. 5: Effects of GSSSG and GSSG on T-cell activation induced by CD3 stimulation or CD3/CD28 co-stimulation.**

**a, b**, FCM analysis of the surface antigens CD25 (**a**) and CD44 (**b**) on naïve CD4^+^ T cells stimulated with anti-CD3ε and anti-CD28 antibodies for 24 h in the presence or absence of GSSSG.

**c, d,** FCM analysis of CD25 on naïve CD4^+^ T cells stimulated with anti-CD3ε antibody (**c**) or anti-CD3ε and anti-CD28 antibodies (**d**) for 24 h in the presence of indicated dose of GSSSG.

**e-h**, Naïve CD4^+^ T cells obtained from WT and *Cars2*^+/–^ mice were stimulated with anti-CD3ε antibodies (**e** and **f**) or anti-CD3ε and anti-CD28 antibodies (**g** and **h**) for 24 h in the presence or absence of GSSG. FCM analysis of the surface antigens CD25 (**e** and **g**) and CD44 (**f** and **h**) on CD4^+^ T cells was performed.

Data are mean ± SD. *P* values were analysed with one-way ANOVA with Tukey’s test. Data represent at least two independent experiments with consistent results. ****P* < 0.001.

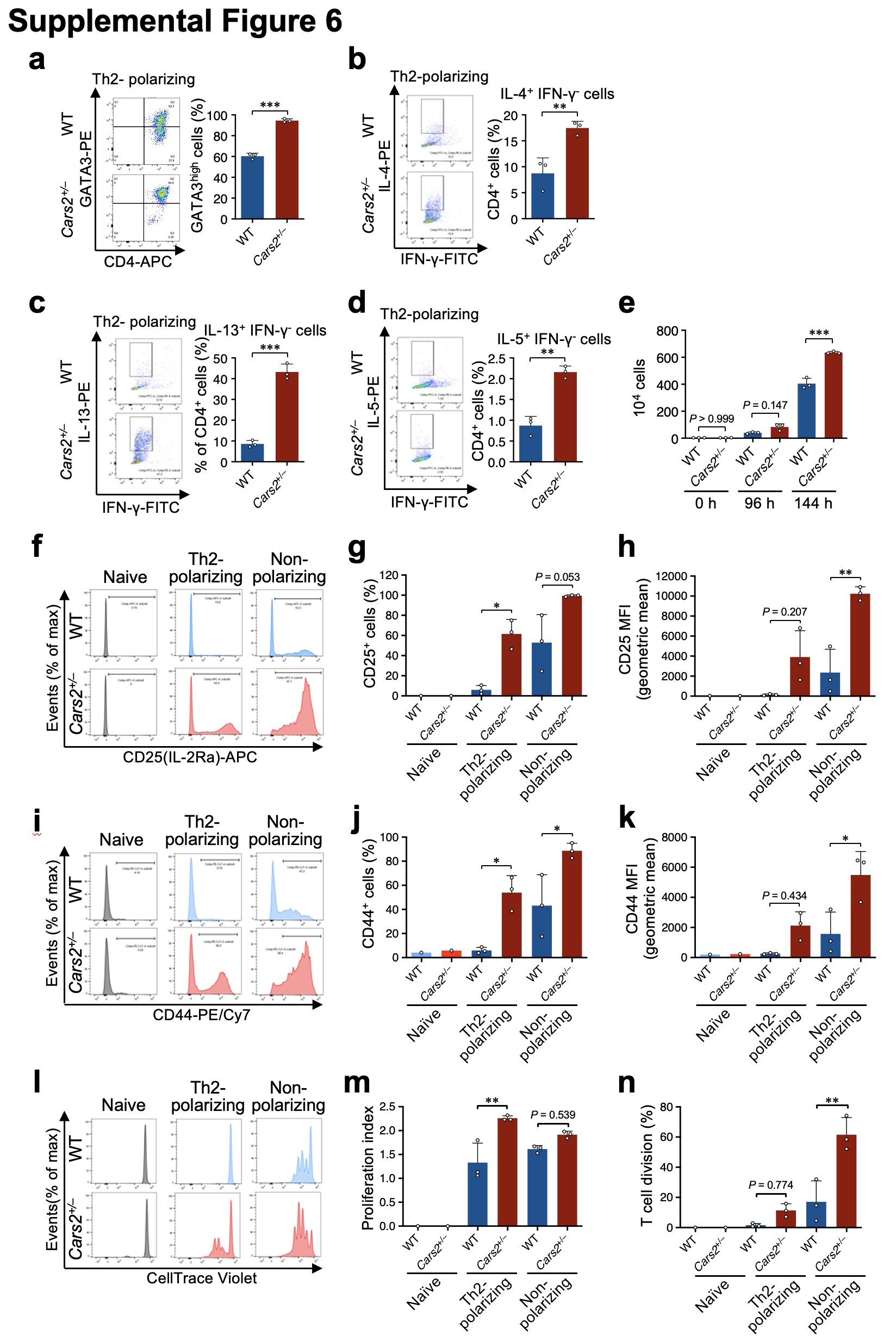

**Supplementary Fig. 6: Enhanced Th2 cell differentiation and proliferation in naïve CD4^+^ T cells from *Cars2*^+/–^ mice.**

**a-d,** Th2 differentiation of naïve CD4^+^ T cells obtained from spleens of WT and *Cars2*^+/–^ mice as analysed by FCM. Representative FCM plots and intracellular expression of GATA3 in CD4^+^ T cells (**a**), and representative FCM plots and intracellular expression of IL-4 (**b**), IL-13 (**c**), and IL-5 (**d**) in CD4^+^ T cells.

**e**, Quantitative analysis of the total numbers of purified T cells cultured under Th2-polarizing conditions for 144 h.

**f-k,** FCM analysis of surface antigens CD25 and CD44 on naïve CD4^+^ T cells after culture under Th2-polarizing conditions or non-polarizing conditions for 72 h. Representative FCM plots (**f**) and surface expression of CD25 in CD4^+^ T cells (**g** and **h**), and representative FCM plots (**i**) and surface expression of CD44 in CD4^+^ T cells (**j** and **k**). **l-n**, CD4^+^ T-cell division analysed by the dye dilution method with CellTrace Violet. Representative FCM plots (**l**), proliferation index (**m**), and percent T-cell division (**n**) determined with FlowJo software.

Data are mean ± SD. *P* values were analysed with a two-tailed Student’s *t*-test (**a-d**) and one-way ANOVA with Tukey’s test (**e**, **g**, **h**, **j**, **k**, **m**, and **n**). Data represent at least two independent experiments. **P* < 0.05; ***P* < 0.01; ****P* < 0.001.

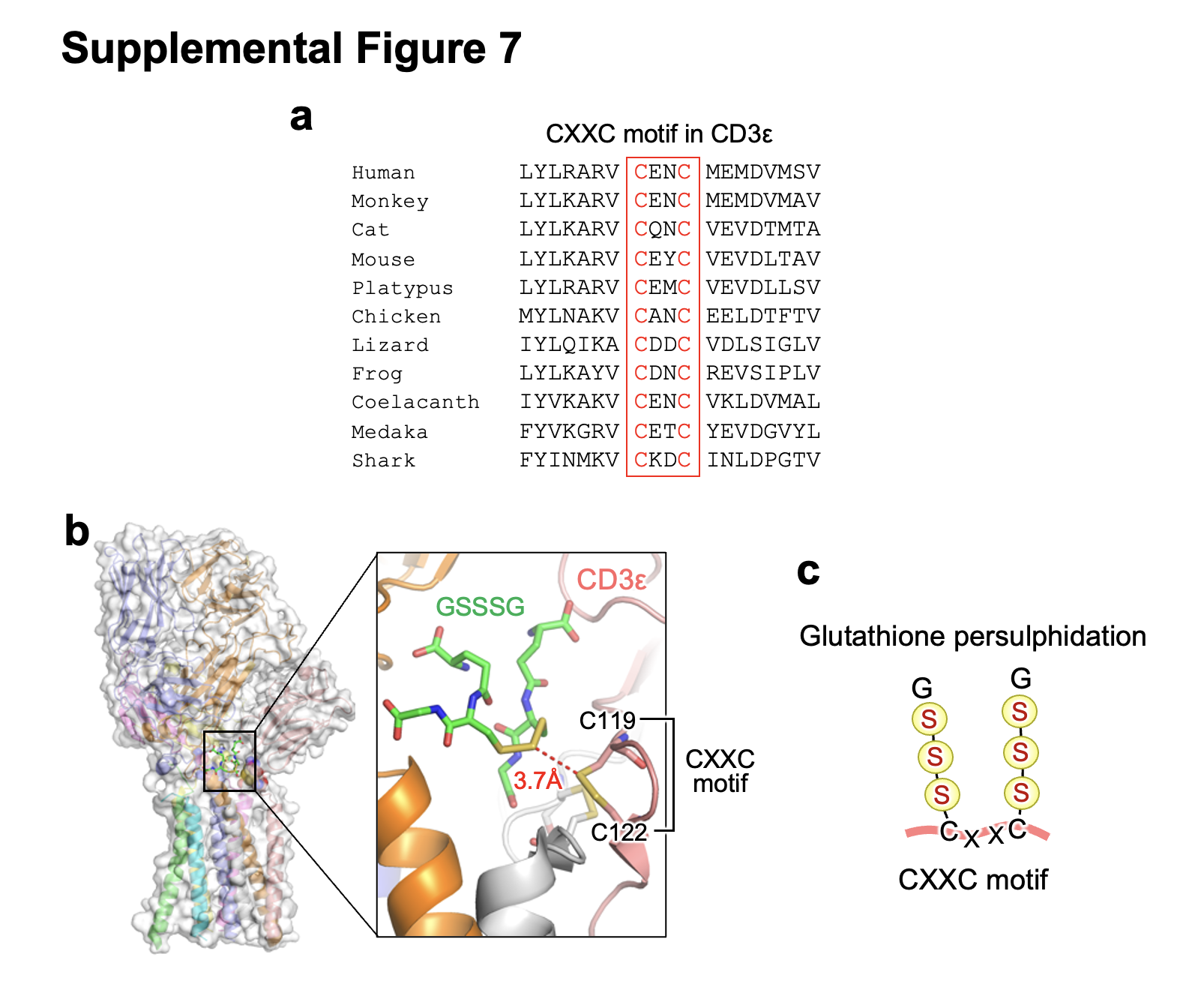

**Supplementary Fig. 7: Persulphidated glutathionylation to the CXXC motif of CD3ε.**

**a,** CXXC motifs in CD3ε chains from various species. Conserved cysteine residues are marked in red.

**b**, Docking simulation showing that GSSSG serves as a ligand for the CD3ε chain in the TCR/CD3 complex. The sulphur atoms of GSSSG are predicted to gain access to the CXXC motif (C119/X/X/C122) of the CD3ε chain; an enlarged view of the area in the rectangle is on the right, where the sulphur atoms of GSSSG and the CXXC motif in CD3ε are represented by yellow sticks.

**c**, Illustration of glutathione persulphide adducts to cysteine residues of the CXXC motif.

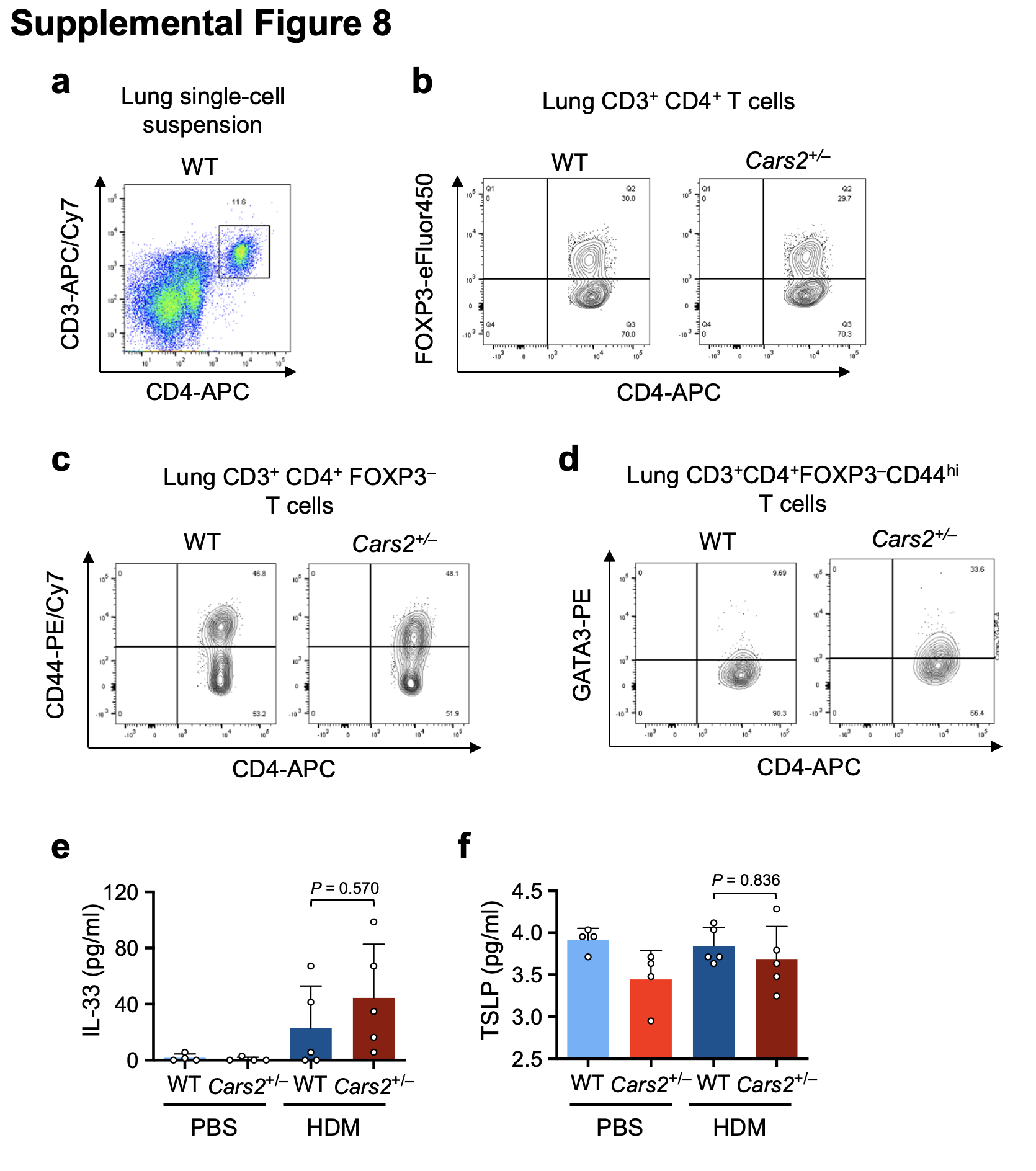

**Supplementary Fig. 8: Enhanced accumulation of GATA3^high^ Th2 cells in the lungs of** ***Cars2*^+/–^ mice.**

**a-d,** Staining strategy to detect GATA3^high^ Th2 cells (CD3^+^CD4^+^FOXP3^–^CD44^hi^ GATA3^high^ cells). The lung cells from WT and *Cars2*^+/–^ mice treated with HDM were stained with antibodies for CD3, CD4 (**a**), FOXP3 (**b**), CD44 (**c**) and GATA3 (**d**). Representative FCM plots were shown.

**e, f,** Type 2 cytokine levels in BALF of a murine asthma model. BALF samples were obtained from the murine asthma model as illustrated in Fig. 5a. Amounts of IL-33 (**e**) and TSLP (**f**) in BALF were measured by using ELISA. Each dot represents data from an individual mouse (*n* = 4–5 per group). Data are mean ± SD. *P* values were analysed with one-way ANOVA with Tukey’s test.

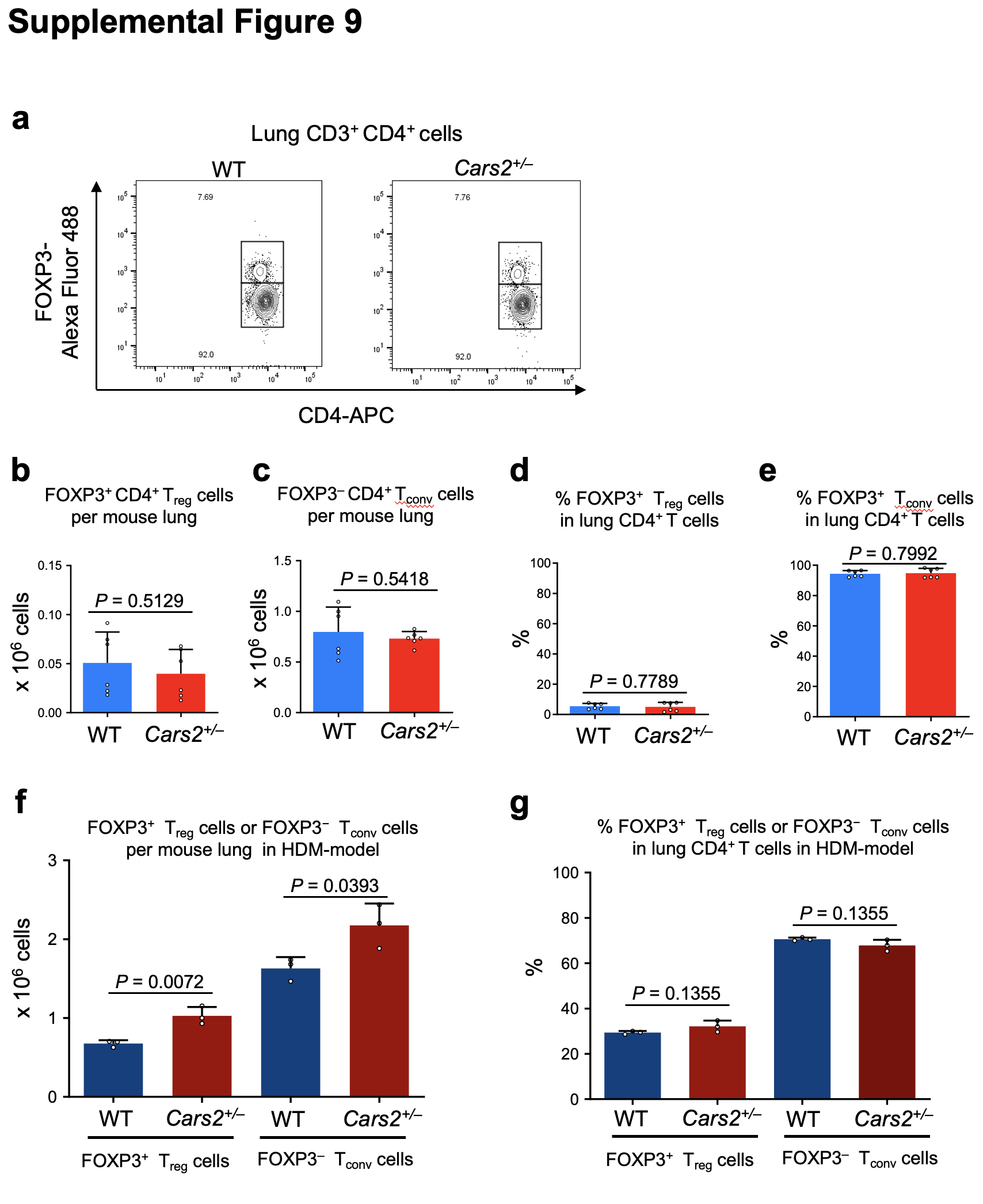

**Supplementary Fig. 9: FCM analysis of lung regulatory T cells from WT and *Cars2*^+/–^ mice at steady state and after HDM-challenge.**

**a,** Representative FCM plots of staining with antibodies for FOXP3 and CD4 in lung CD4^+^ T cells obtained from WT and *Cars2*^+/–^ mice at steady state.

**b-e,** The number of lung regulatory T (Treg) cells (**b**) and conventional CD4^+^ T (Tconv) cells (**c**), and the percent of Treg cells (**d**) and Tconv cells (**e**) in total lung CD4^+^ T cells in WT and *Cars2*^+/–^ mice at steady state.

**f, g,** The number of Treg cells and conventional Tconv cells per mouse lung (**f**), and the percentage of Treg cells and Tconv cells in total lung CD4^+^ T cells (**g**) in WT and *Cars2*^+/–^ mice treated with HDM.

Data are mean ± SD. *P* values were analysed with Student's *t*-test (**b-g**).

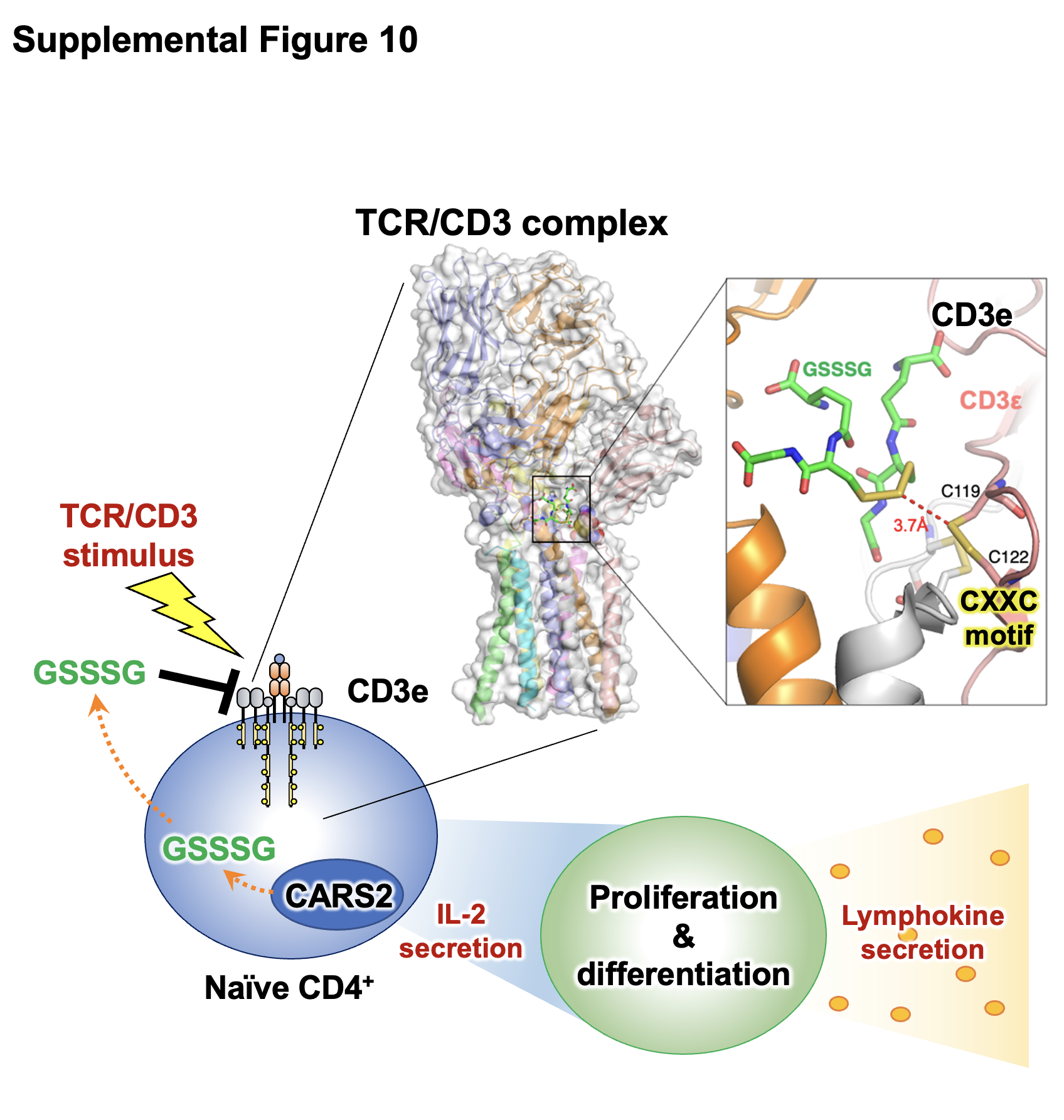

**Supplementary Fig. 10: Schematic illustration of supersulphide-mediated regulation of TCR signalling.**

CARS2 produces supersulphides including GSSSG, which is a major endogenous supersulphides. GSSSG likely suppresses TCR signal activation by conjugating to the CXXC motif of the CD3ε chain in the TCR/CD3 complex, resulting in the suppression of T-cell proliferation and differentiation in response to TCR/CD3 stimulation. Considering the downregulation of Cars2 in naïve CD4^+^ T cells in response to TCR stimuli, the CD3ε chain in the basal state is likely to undergo persulphidated glutathionylation, which is interpreted as an active suppression of T cells to avoid their aberrant activation. Supersuphide reduction in T cells upon stimuli is thus interpreted as a reasonable response to allow their full activation. From a therapeutic point of view, GSSSG is a promising anti-inflammatory agent for allergic and autoimmune diseases.
